# Interplay of condensation and chromatin binding underlies BRD4 targeting

**DOI:** 10.1101/2024.02.07.579384

**Authors:** Amy R. Strom, Jorine M. Eeftens, Yury Polyachenko, Claire J. Weaver, Hans-Frederick Watanabe, Dan Bracha, Natalia D. Orlovsky, Chanelle C. Jumper, William M. Jacobs, Clifford P. Brangwynne

## Abstract

Nuclear compartments form via biomolecular phase separation, mediated through multivalent properties of biomolecules concentrated within condensates. Certain compartments are associated with specific chromatin regions, including transcriptional initiation condensates, which are composed of transcription factors and transcriptional machinery, and form at acetylated regions including enhancer and promoter loci. While protein self-interactions, especially within low-complexity and intrinsically disordered regions, are known to mediate condensation, the role of substrate-binding interactions in regulating the formation and function of biomolecular condensates is under-explored. Here, utilizing live-cell experiments in parallel with coarse-grained simulations, we investigate how chromatin interaction of the transcription factor BRD4 modulates its condensate formation. We find that both kinetic and thermodynamic properties of BRD4 condensation are affected by chromatin binding: nucleation rate is sensitive to BRD4-chromatin interactions, providing an explanation for the selective formation of BRD4 condensates at acetylated chromatin regions, and thermodynamically, multivalent acetylated chromatin sites provide a platform for BRD4 clustering below the concentration required for off-chromatin condensation. This provides a molecular and physical explanation of the relationship between nuclear condensates and epigenetically modified chromatin that results in their mutual spatiotemporal regulation, suggesting that epigenetic modulation is an important mechanism by which the cell targets transcriptional condensates to specific chromatin loci.

## Introduction

Phase separation is a process by which molecules within living cells can be concentrated into biomolecular condensates, membrane-less compartments that promote enzymatic activities at targeted sites^1,2^. Within the nucleus, multiple condensates exist that organize specialized functions within the complex nuclear environment, including ribosome production within nucleoli^3,4^, mRNA splicing within nuclear speckles^5–8^, as well as chromatin remodeling and transcriptional initiation within transcription factor condensates^9,10,11–14^. Formation of these nuclear condensates at specific genomic loci is essential for their function: nucleoli form at rDNA^3,4^, nuclear speckles associate with highly transcribed genes^5–8^, and localized condensation of chromatin remodelers and transcription factors dictates gene expression^9^. Still, both experimental and simulation studies of the biophysical determinants of condensation have, to date, focused primarily on describing interactions that drive condensate formation, leaving the mechanisms underlying their genomic targeting relatively unstudied. Transcription factors often contain dual modalities: both self-interaction motifs to promote condensation, and chromatin-binding motifs to interact with certain genomic regions, though the biophysical interplay of these multiple types of interactions and their potential to drive localized condensation have not been fully explored.

Condensation is driven by thermodynamic demixing of biomolecules due to transient multivalent interactions, which are often provided by intrinsically disordered regions of proteins (IDRs), though multivalent interactions can also arise from folded protein regions or from binding to substrates including RNA or chromatin^15^. When the concentration of a phase-separation-prone protein is above a threshold called the saturation concentration, it becomes energetically favorable for the solution to phase separate, forming a continuous dilute and a discrete dense phase of droplets^16^. Phase separation-prone proteins including transcription factors often incorporate both homotypic (self-self) and heterotypic (self-nonself) interactions, both of which may modify the condensation properties of the phase separating system^17,18^.

Targeted condensation can occur through seeded nucleation on a substrate, a process that is driven by kinetic properties of the phase separating system. Classical nucleation theory describes the initial formation of a condensed phase from an homogenous solution, which occurs when freely diffusing monomers begin to gather into dynamic nanoscale clusters^19^. These clusters continually form and dissolve, until one exceeds the critical radius, at which point it is more energetically favorable to grow than shrink. Many features of the nucleation of biomolecular condensates are in agreement with classical nucleation theory, including the strong dependence of nucleation rate on the degree of supersaturation^20–22^. Targeted nucleation can be achieved by lowering the energetic barrier of nucleation through binding monomers to a substrate or ‘seed,’ which reduces the critical radius required for continued growth, in a process referred to as heterogeneous nucleation^19,23^. Simulations have demonstrated that seeded nucleation results in faster and spatially localized condensate formation, and may be an important mechanism through which functional endogenous condensates are targeted to form at specific genomic loci^20,21,24^, though this has not yet been experimentally demonstrated.

Transcription factor condensates are an ideal model to study the kinetic and thermodynamic contributions of chromatin substrate binding to functional biomolecular phase separation, as the ability of transcription factors to both condense and bind chromatin is well established^11,12,25,26^, and their targeting to specific genomic loci is functionally relevant in dictating the cell’s transcriptional profile. The BET family protein BRD4 is a well-studied transcription factor that localizes to acetylated chromatin sites^27^, recruits pTEF-b^28^ and initiates transcription of key genes involved in signal response, immunity and oncogenesis^29^. All BET family proteins (BRD2, BRD3, BRD4 and BRDT) bind to lysine-acetylated histones through two conserved N-terminal bromodomains, and recruit transcriptional initiating cofactors through the shared ET domain. However, only BRD4 can directly engage pTEF-b through a C-terminal motif (CTM) in its extended, unstructured C-terminus. Moreover, while the expression of BRD2, BRD3 and BRDT are tissue-specific, BRD4 expression is ubiquitous^29^. BRD4’s C-terminal ∼1000 amino acid intrinsically disordered region (IDR) has been characterized as a driver of self-interaction and condensate formation in living cells^12,30–32^, which has been implicated in its enhanced ability to drive high levels of transcription.

For BRD4 specifically, condensate-forming capabilities of the C-terminal IDR are thought to occur through self-interactions, or in combination with MED1^12^, which contributes to the thermodynamic driving force for phase separation. Localization to specific chromatin regions is mediated through the two N-terminal bromodomains, which bind acetylated histones and thereby serve as reader domains to mediate genomic interaction. The inherent multivalency of potential acetylation regions on the genome may not only provide targeting sites that can enhance kinetics, but could also potentially contribute thermodynamically to BRD4 phase separation^12,33^. Indeed, substrates with repetitive binding sites may enhance the valence of the condensing system and allow formation of condensates at lower concentration. This is seen with other nucleic acid-binding proteins like FUS, which is capable of condensation without RNA-binding, while addition of RNA reduces the critical concentration^34,35^. However, it remains to be elucidated how the interplay of kinetic and thermodynamic properties of transcription factor condensates facilitate their formation at targeted chromatin loci.

Here we use coarse-grained simulation and quantitative experiments in living cells to decode the biophysical rules underlying how interactions between chromatin and proteinaceous condensates of BRD4 impact condensate formation and targeting. We show that binding to acetylated chromatin regions allows BRD4 condensation at lower valence than is required for off-chromatin, and that these acetylated regions act as heterogeneous nucleation seeds for BRD4 condensates. We propose that the cell regulates both kinetic and thermodynamic properties of condensation to control the spatiotemporal localization of transcriptional condensates for their nuclear function.

### Disruption of BRD4 chromatin binding reduces number and volume of condensates

The protein structure of BRD4 includes two folded N-terminal bromodomains (BD1 aa 75-247; BD2 aa 360-440) which bind to acetylated chromatin, and a C-terminal motif (CTM aa 1047-1362) that interacts with pTEF-b, as well as the large C-terminal IDR which coordinates self-interaction and is known to contribute to BRD4 phase separation^12,31^ (Fig. 1A). By immunofluorescence, and in agreement with previous reports^12,13^, BRD4 forms punctate nuclear structures under endogenous expression levels in U2OS cells, which are recapitulated with expression of fluorescently-tagged BRD4^FL^-mCherry in living nuclei (Fig. 1B). To examine the role of chromatin binding in BRD4 condensation and targeting, we utilized the small molecule BET inhibitor (BETi) JQ1, which can specifically disrupt interaction of bromodomains with acetylated chromatin^36^. Upon treatment of cells with 1 μM JQ1 for 90 min, we find that BRD4 becomes dispersed throughout the nucleoplasm, with few large foci remaining per nucleus in both the endogenous immunofluorescence and exogenous expression cases (Fig. 1B). This altered nuclear localization is phenocopied by expression of the truncated construct BRD4^ΔN^-mCherry (aa 441-1362) in which both bromodomains are removed (Fig. 1A, B). A panel of cells arranged by expression level of BRD4^FL^-mCherry shows that cells with higher expression have more puncta (Fig. 1C). Nonetheless, across expression levels the number of puncta in individual cells after JQ1 treatment is consistently lower than before treatment, visually resembling cells expressing BRD4^ΔN^-mCherry at similar expression levels (Fig. 1C). To avoid BRD4 expression-dependent effects in quantification, we used the number and size of endogenous BRD4 puncta by immunofluorescence to create lower and upper gates on the relevant expression levels of BRD4^FL^-mCherry (Fig. 1C), such that within this range, the quantification of the number and size of live expression BRD4^FL^-mCherry condensates is indistinguishable from quantification of the endogenous protein by immunofluorescence. This analysis further confirms that disruption of chromatin binding with JQ1 results in fewer, larger puncta that match those formed by BRD4^ΔN^-mCherry (Fig. 1D).

**Figure 1:**
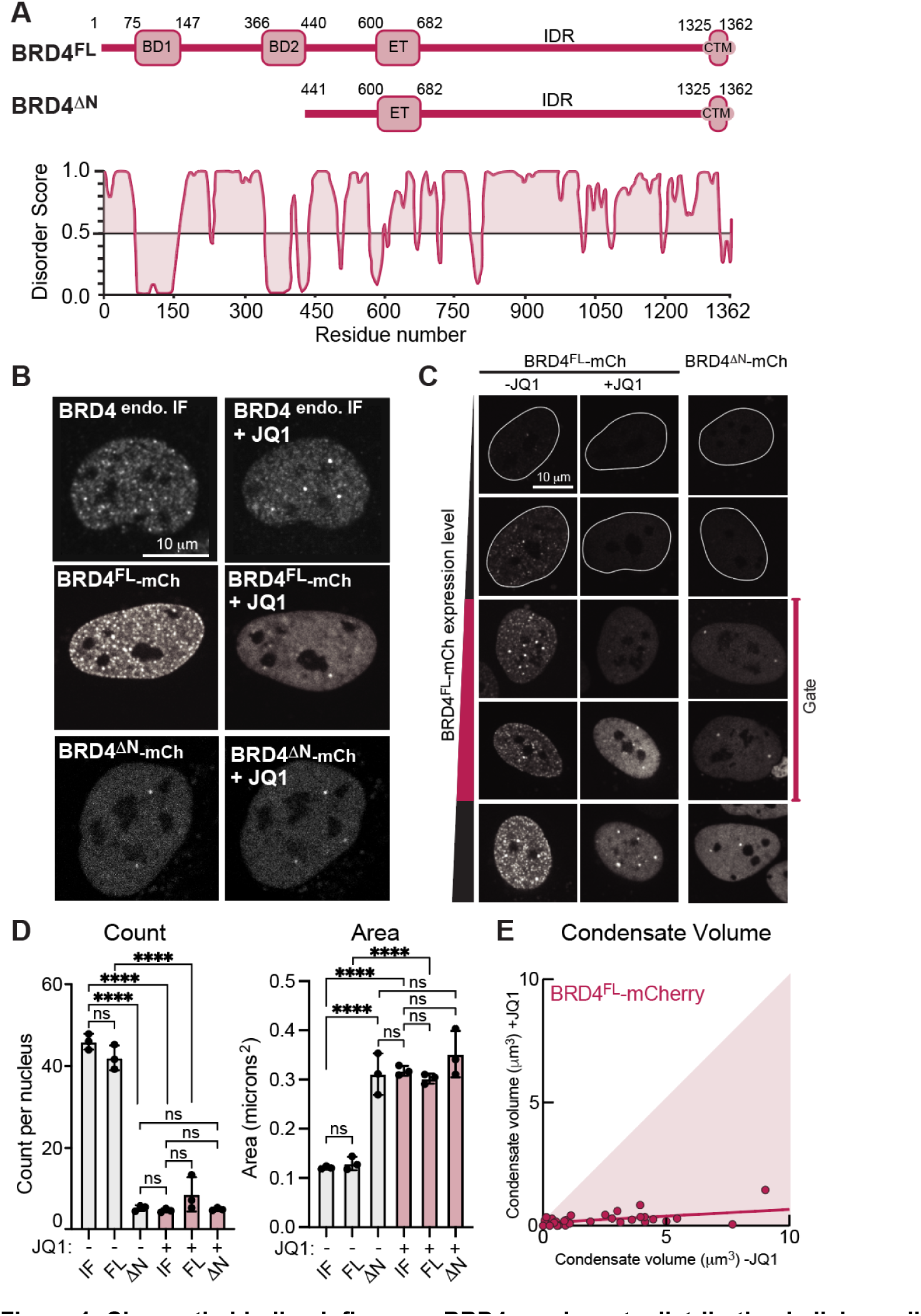
Chromatin binding influences BRD4 condensate distribution in living cells. **A.** Schematic of BRD4 full length (FL) and deltaN-terminus (ΔN) constructs, as well as PONDR predicted disorder score. **B.** From top: Immunofluorescence of endogenous BRD4 protein in cultured human U2OS cells without and with addition of 1 µM JQ1. Exogenous live expression of BRD4^FL^-mCherry and BRD4^ΔN^-mCherry in the same cell before (-JQ1) and after (+JQ1) addition of 1 µM JQ1. **C.** Panel of images of U2OS cells with increasing expression level of BRD4^FL^-mCherry (same cell -/+ JQ1) or BRD4^ΔN^-mCherry at similar expression levels. **D.** Quantification of number and size of BRD4 condensates from immunofluorescence of endogenous, or live expression of BRD4^FL^-mCherry or BRD4^ΔN^-mCherry, -/+ JQ1. Points represent averages of 3 biological replicates of 25 cells each within the expression level gate defined in 1C, error bars S.E.M. Statistical test One-way ANOVA, ****p < 0.0001. **E.** Estimated condensate Volume measured in the same set of cells expressing BRD4^FL^-mCherry before (x-axis) and after (y-axis) disruption of chromatin binding through addition of 1 µM JQ1. If condensate volume is not affected, points should lie on the diagonal, indicated by shading.

These findings demonstrate that the number, size and positioning of BRD4 condensates in the nucleus is altered by chromatin binding capabilities. A simple explanation is that disruption of chromatin binding releases condensates from defined loci, allowing them to coalesce into fewer, larger droplets^37–39^. If the alteration of condensate number and size is purely due to kinetic effects, we would expect the total volume of condensed material in each cell to remain consistent before and after addition of JQ1, such that a set volume of many small droplets reorganizes into fewer, larger ones. To test this, we measured the cross-sectional area of BRD4^FL^-mCherry condensates in the same nuclei before and after JQ1 treatment, then estimated the volume of condensates in each nucleus (see Methods). Surprisingly, we find that the volume of BRD4^FL^-mCherry condensates is substantially decreased upon JQ1 treatment (Fig. 1E), suggesting that disruption of BRD4’s chromatin binding does not simply lead to coalescence of small droplets, but also impacts the thermodynamics of BRD4 condensation. Together, these data illustrate a complex role for chromatin binding in regulating both kinetic and thermodynamic properties of BRD4 condensation that warrants further quantitative investigation.

### Chromatin binding thermodynamically enhances BRD4 condensation

We next sought an experimental system that would be capable of quantitatively investigating the concentration-dependent thermodynamics of BRD4 phase separation, while also having a well-defined trigger for initiating condensate formation to study kinetics. The Corelet system is a two-component platform capable of triggering phase separation in living cells through light-induced oligomerization of an sspB-tagged phase-separation-prone protein domain via interaction with 24-mer multivalent iLID-GFP-Ferritin ‘Cores’^40^. The Corelet system can be used to build a phase diagram to map the thermodynamic properties of BRD4 phase separation by quantitatively determining the Core concentration and BRD4-to-Core valence required for BRD4^FL^ condensate formation in untreated (Control) or JQ1-treated cells (with disrupted chromatin binding).

We expressed BRD4^FL^ Corelet components in U2OS cells and initiated their oligomerization with blue light (Fig. 2A). Before light activation, live expression of BRD4^FL^-mCherry-sspB exhibits a few puncta pre-activation, similar to the native condensates visualized by immunofluorescence (Fig. 2A, left). Then, upon light-activated oligomerization, de novo BRD4^FL^ condensates form (Fig. 2A, right). Because oligomerization in the Corelet system is reversed in the absence of blue light, we can compare the number and size of condensates in the same set of BRD4^FL^-mCh-sspB expressing cells before and after addition of JQ1 (1μM, 90 min), to determine if the concentration and valence required for condensation are altered upon loss of chromatin binding. In the same cell with JQ1 treatment, light activation again leads to BRD4^FL^ Corelet condensate formation (Fig. 2B), though the condensates are fewer and larger (Fig. 2C), mimicking the changes observed upon disruption of chromatin binding in the endogenous and exogenous expression systems introduced earlier. JQ1 treatment of BRD4^ΔN^ Corelet condensates has no effect on puncta number or size (Fig. 2D, Fig. S1A-D), confirming that JQ1 specifically disrupts chromatin binding, without affecting IDR-mediated condensate formation. The number of Corelet puncta per nucleus on a cell-by-cell basis before and after addition of JQ1 makes clear the drastic reduction in BRD4^FL^-mCh-sspB condensate number at most expression levels (Fig. S1D). Interestingly, cells with very high Corelet condensate count are less severely affected, likely because oligomerization through the Ferritin Core platform can substitute for oligomerization through chromatin-binding at very high expression levels, making these cells less responsive to JQ1.

**Figure 2.**
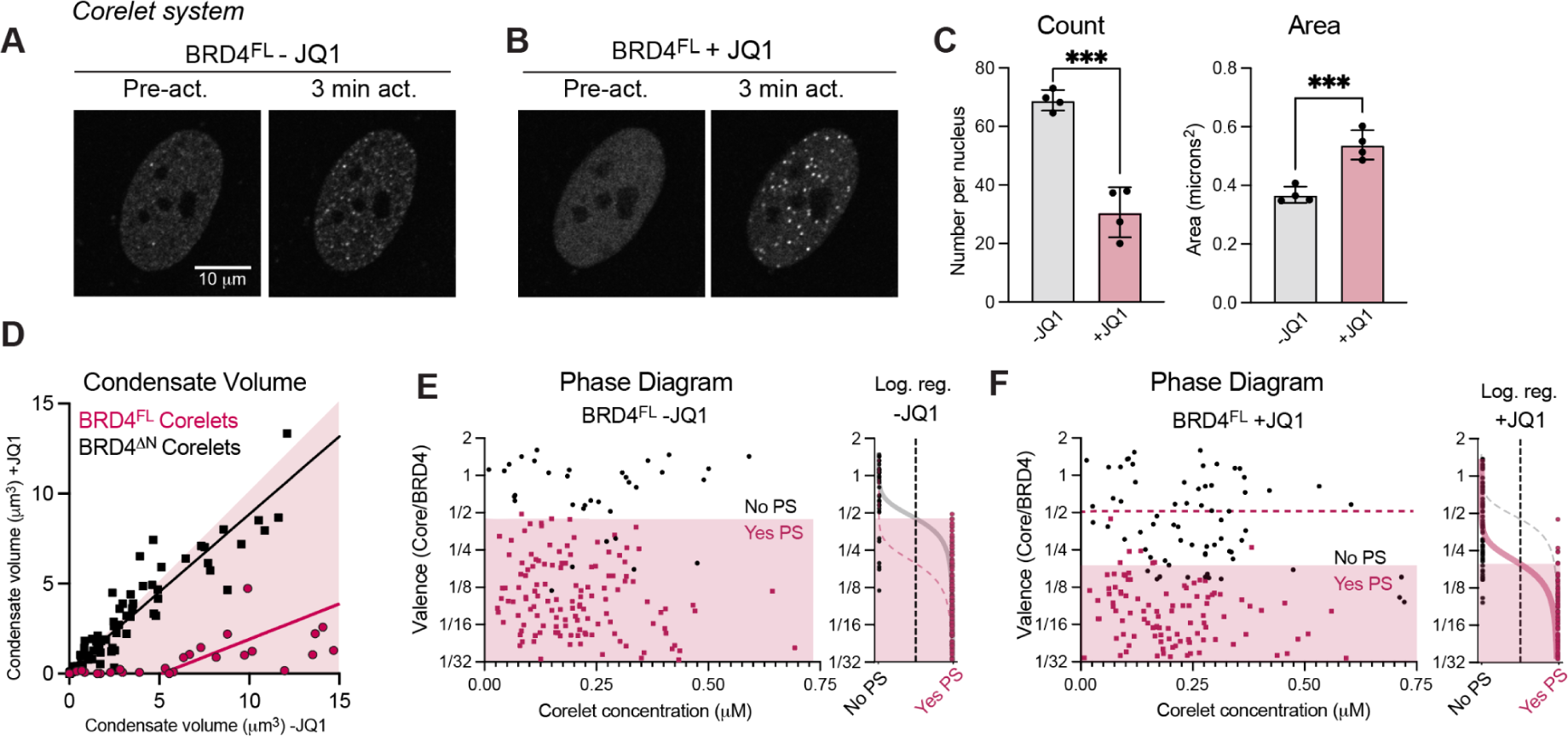
Thermodynamic effects of BRD4 chromatin binding are measured the Corelet synthetic oligomerization system. Representative images of BRD4^FL^ Corelets pre- and post-light-activation of a nucleus without **(A)** or with **(B)** 1µM JQ1. **C.** Quantification of the number and size of BRD4^FL^ Corelet condensates per nucleus induced with light activation -/+ JQ1 in the same set of nuclei. Error bars represent SEM across four trials of 25 cells each, Students t-test ***p = 0.0002 in Count, *** p= 0.001 in Area. **D.** Condensate Volume comparison in -/+JQ1 conditions in the same set of cells with BRD4^FL^ (pink) or BRD4^ΔN^ (black) Corelets. BRD4^ΔN^ Corelet condensate volume is unaffected by the addition of JQ1, as can be seen from points largely along the diagonal (shaded pink), while BRD4^FL^ Corelet condensate volume is much lower after addition of JQ1. Lines are linear fit. **E-F.** Phase diagram and logistic regression of BRD4^FL^ measured with the Corelet system in the absence **(E)** and presence **(F)** of JQ1. Phase diagrams are constructed from the same cells expressing BRD4FL -/+ JQ1 demonstrating the shifted valence between -JQ1 (gray) and +JQ1 (pink).

Calibrated fluorescence microscopy of cells expressing BRD4^FL^ Corelets can be used to construct an oligomerization- and concentration-dependent phase diagram, wherein proteins that form condensates at lower BRD4-to-Core valence have a stronger phase separation tendency (plotted as inverse valence, Core/BRD4). Compared to previously published examples of Corelet phase diagrams^40,41^, extremely low Core concentration is sufficient to initiate BRD4^FL^ condensate formation, and we measure a critical valence of two BRD4^FL^-mCh-sspB bound per core required for condensate formation (shaded area, Fig. 2E). This lack of a left-hand boundary at low Core concentration highlights that the nuclear interior is poised for BRD4 phase separation with minimal additional oligomerization and may reflect the presence of other interaction partners in the nucleus like MED1^12^. The presence of BRD4 interactors like MED1 in the nucleus could contribute to the pre-existing puncta seen before light activation and make the saturation concentration appear lower than it would be for BRD4 alone. When the valence distinction between phase-separating (Yes PS) and non-phase-separating cells (No PS) is measured in the same nuclei after disruption of chromatin binding with JQ1, it is shifted to five BRD4^FL^-mCh-sspB bound per core required for condensate formation (Fig. 2F). This shift in required valence is readily apparent when the phase diagram data is quantified as a logistic regression (Fig. 2E, F). Together, these data indicate that BRD4 is capable of condensate formation in the Corelet system both with and without chromatin binding, but that chromatin binding provides a thermodynamic enhancement of BRD4’s phase separation tendency.

**Figure S1.**
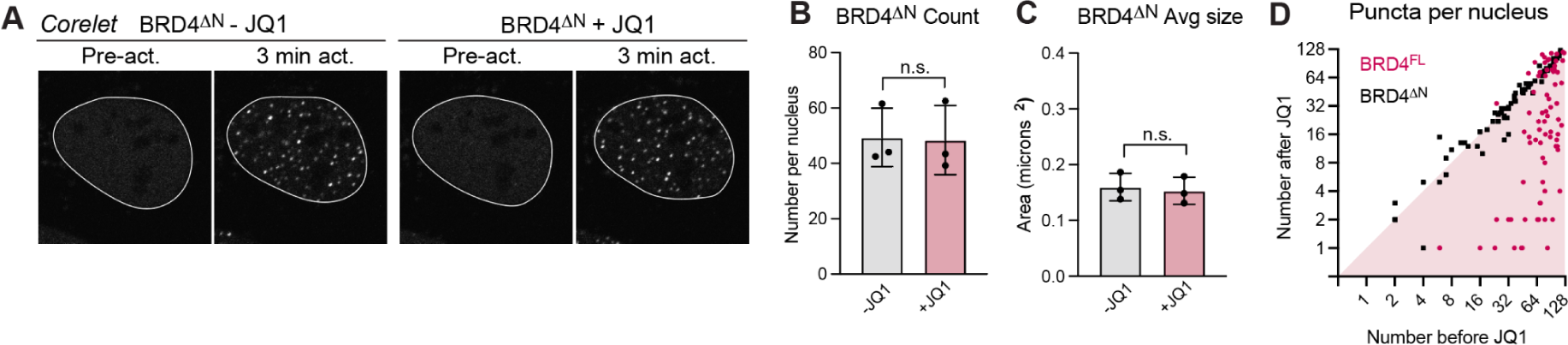
JQ1 does not affect BRD4^ΔN^ puncta, related to Figure 2. **A.** Representative images of BRD4^ΔN^ light-induced Corelet-condensates +/- JQ1 treatment. Quantification of count (**B**) and area (**C**) of BRD4^ΔN^ Corelet condensates +/- JQ1 in 3 trials of 25 cells each. Student’s t-test. **D.** Single-cell BRD4^FL^ Corelet-condensate count per nucleus (pink) before JQ1 treatment (X-axis) and after JQ1 treatment (Y axis). Each point is one nucleus. Number of BRD4^ΔN^ Corelet condensates per nucleus (black) are unaffected by JQ1 and lie on the diagonal.

### Coarse-grained simulations characterize valence-dependence of BRD4 condensation

To better understand the thermodynamic and kinetic mechanisms by which chromatin binding affects BRD4 phase separation, we next created a coarse-grained simulation of BRD4 condensation. In this system (Fig. 3A), the chromatin polymer is represented by a chain of nucleosome monomers, each with eight histone tails that can either be in an acetylated state, enabling attractive interactions with BRD4 (green), or in a non-interacting state (red). The ∼1400 amino acid BRD4 protein is represented as two soft spheres^42^ connected by a harmonic potential, one representing the acetylated histone tail-interacting N-terminus (referred to as “chromatin-interacting” in what follows) and the other representing the disordered self-interacting C-terminus. Model BRD4 Corelets of variable valence are created by tethering BRD4 molecules to a central oligomerization platform via their C-termini, mimicking the geometry of these constructs in the Corelet system.

**Figure 3.**
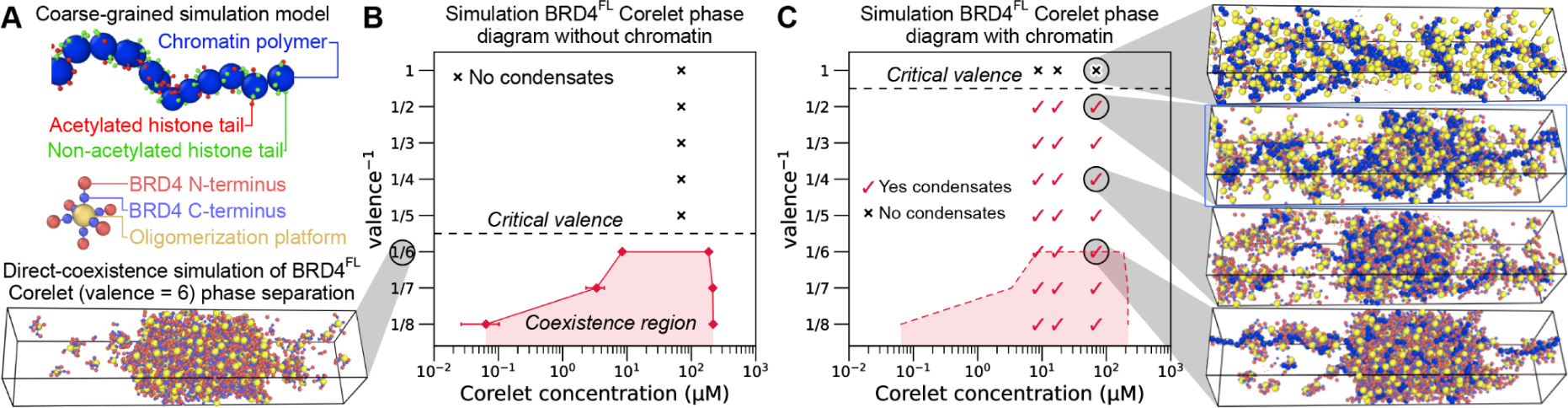
A coarse-grained model for simulating BRD4 Corelet condensation. **A.** Coarse-grained simulations contain representations of the chromatin polymer (blue) and Corelet oligomerization platform (gold) with attached BRD4 molecules, each composed of two spheres representing the N-terminus (capable of interacting with chromatin) and C-terminus (capable of self-interaction). **B.** The valence-dependent phase diagram in the absence of chromatin, obtained via direct-coexistence simulation (representative snapshot on left), shows a critical valence^-1^ of 1/6. **C.** A phase diagram showing the presence of chromatin-Corelet condensates in simulations with strong chromatin binding (40% acetylated histone tails) predicts an apparent critical valence^-1^ of 1/2. Representative snapshots are shown (on right) for the indicated simulation conditions.

Simulations of model BRD4 Corelets can be used to create phase diagrams that parallel the experimental Corelet phase diagrams from Figure 2, both in the presence and absence of interactions with acetylated chromatin. Direct-coexistence simulations^43^ of model Corelets indicate that the critical valence for phase separation in the absence of chromatin is between 5 and 6 BRD4 molecules per ‘Core’ oligomerization platform (Fig. 3B), which is in close agreement with the experimental measurement of critical valence from Figure 2F. We note that a key difference between the simulation and experimental phase diagrams is that all simulations are performed with a monodisperse valence distribution, whereas Corelets in experiments likely have a broader valence distribution. Broadening the valence distribution tends to reduce the apparent saturation concentration for phase separation, since higher-valent species tend to phase-separate at lower concentrations^44^. When chromatin with 40% acetylated histone tails is added, this results in the formation of chromatin-enriched condensates at low valence, in conditions where BRD4 Corelets are unable to condense on their own (Fig. 3C). These simulations indicate that the critical valence is two BRD4 molecules per oligomerization platform in the presence of acetylated chromatin, due to a combination of chromatin-BRD4 and BRD4-BRD4 attractive interactions^45,46^. This is again in striking agreement with experimental findings (Fig. 2E).

Importantly, our simulations provide an explanation for the critical valence difference between the -/+ JQ1 cases in experiments, since the elimination of chromatin-BRD4 interactions reverts the phase behavior to that of the Corelet system without chromatin. Moreover, the simulations suggest that puncta visualized at very low valence in the -JQ1 experimental case do not represent conventional phase separation, but rather subsaturation clusters stabilized by regions of highly acetylated chromatin, because the highly acetylated chromatin is a necessary component of the condensate. In our simulations, these clusters do not grow to be much larger than the globule of compacted chromatin. BRD4 clustering in this scenario does not correspond to a unique coexistence pressure, which would determine the saturation concentration of Corelets in conventional phase separation. Consistent with this interpretation, condensates with valence^-1^ between 1/2 and 1/5 are limited in volume (Fig. S2A-B) and entirely dissolve upon disruption of chromatin binding (Fig. S2C). Quantification of the fraction of Corelets in the dense (liquid) phase in simulated condensates (Fig. S2D) are consistent with measured volume of condensates in Corelet experiments (Fig. S2C) and demonstrate that below the critical valence^-1^ (1/6, dotted line in Fig. S2D), we do not observe a consistent dense phase fraction. Together these data indicate that chromatin is an integral part of BRD4 condensates that form at low BRD4-to-Core ratios and at low Corelet concentrations, suggesting that chromatin interactions can affect the thermodynamics of BRD4 condensate formation.

**Figure S2.**
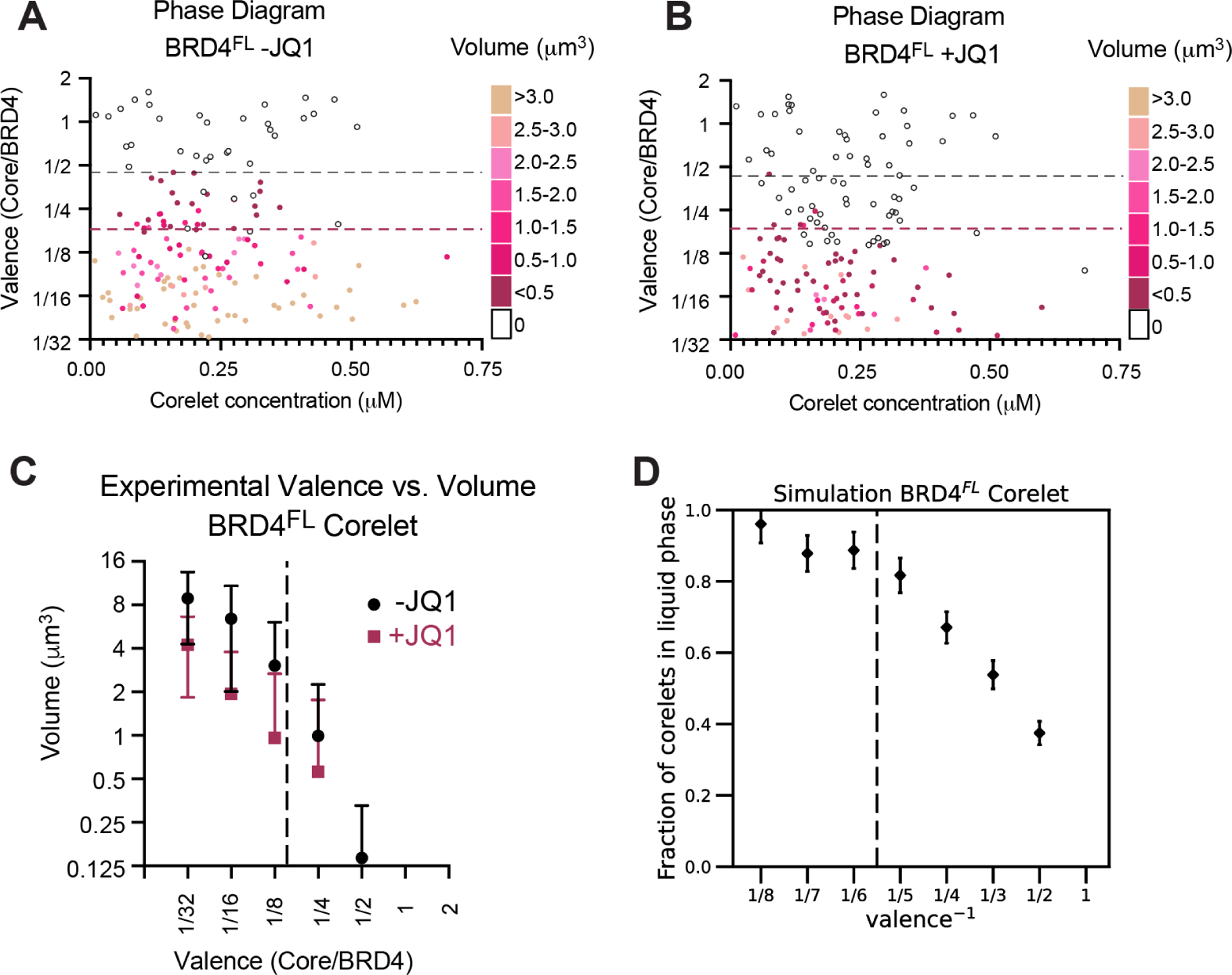
Valence-dependence of BRD4 Corelet condensate volume, related to Figure 3. **A-B.** Experimental phase diagrams -/+ JQ1 recolored by average volume of condensed phase per nucleus. **C.** Phase diagram condensate volume data from -/+JQ1 cells plotted with volume as the independent variable. Note that clusters below valence^-1^ 1/5 are very small, consistent with simulation predictions. (-JQ1 n = 16, 37, 36, 41, 14; +JQ1 n = 16, 36, 37, 12, 0 cells in valence bins 1/32, 1/16, 1/8, 1/4, 1/2, respectively). Error bars SD. **D.** Simulation measurements of fraction of corelets in liquid (dense) phase, demonstrating that below the critical valence (dotted line), clusters can form but no stable dilute fraction is achieved.

### Chromatin acts as a heterogeneous seed for BRD4 condensate nucleation in cells

Because the chromatin-bound clusters of BRD4 exhibit some non-canonical condensation behaviors in phase diagrams, we sought to determine whether classical nucleation theory still applies to BRD4 condensate formation. If chromatin binding acts as a heterogeneous seed for BRD4 condensation, nucleation theory predicts that altered chromatin binding will manifest in changes to the following experimentally-accessible metrics: 1) the rate of condensate formation, 2) the delay time before visible condensates appear, and 3) the steady-state number density (number per unit area) of condensates (Fig. 4A, B). The third outcome arises because the growth of the first condensates to form suppresses the nucleation of additional condensates elsewhere^47,48^, and subsequent coalescence is extremely slow. Leveraging the high spatiotemporal control over condensate initiation enabled by the Corelet system, we measured these three parameters of BRD4^FL^ nucleation before and after addition of JQ1 (1 μm, 90 min). We find that disrupting chromatin binding leads to significantly lower nucleation rates, longer delay times, and ultimately a lower number density of condensates in JQ1 treated cells compared to the same cells before JQ1 treatment (Fig. 4C-E). These observations indicate that chromatin binding aids BRD4^FL^ condensate nucleation in this Corelet system assay.

**Figure 4:**
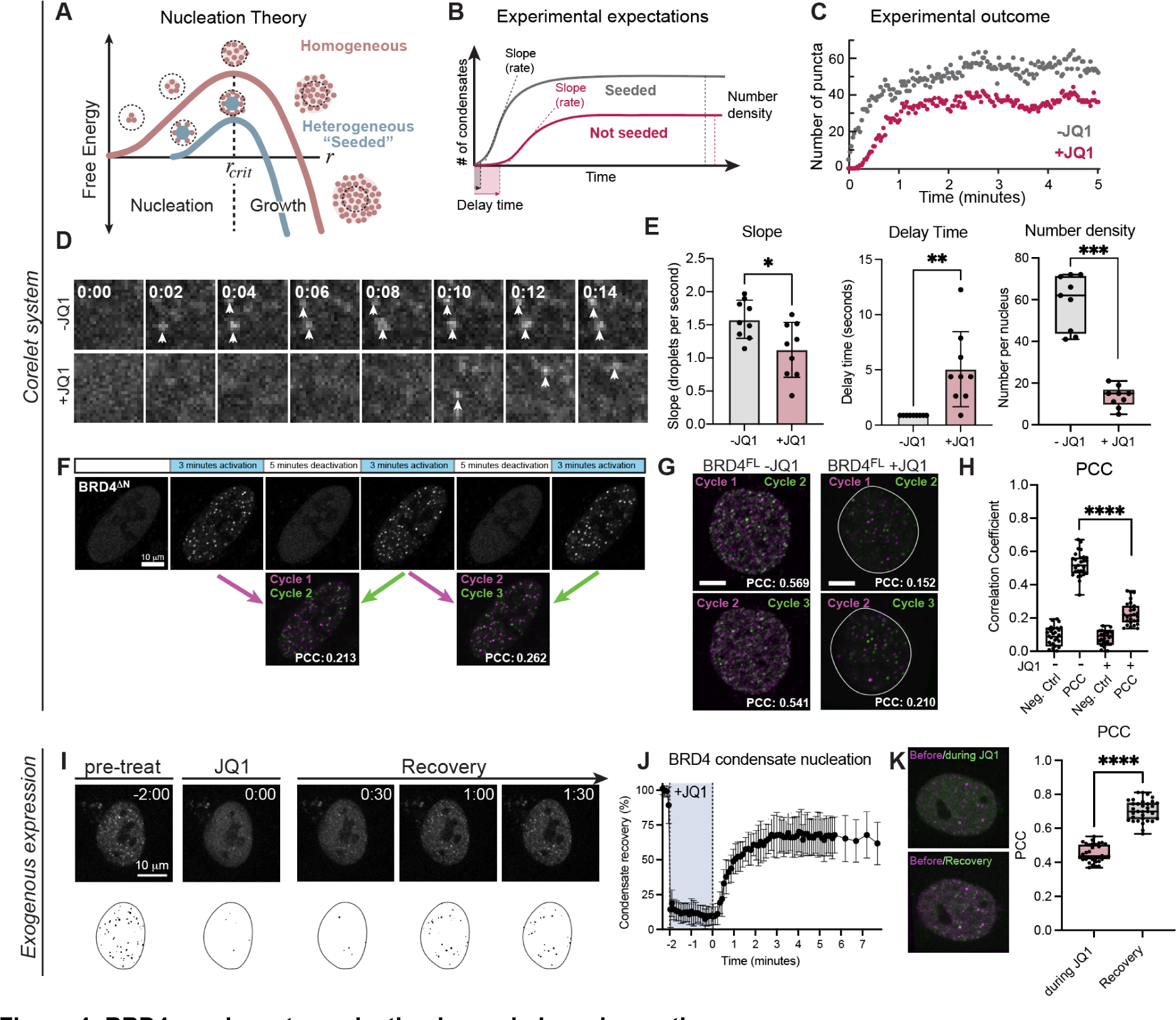
BRD4 condensate nucleation is seeded on chromatin. **A.** Schematic of two modes of condensate nucleation: substrate-seeded (heterogeneous) and not seeded (homogeneous). Without seeding, the free energy of protein clustering increases until the cluster reaches its critical radius (*r_crit_*), at which point it becomes energetically favorable to grow. A seed can lower the energetic barrier to reach *r_crit_*. **B.** Seeded nucleation is expected to have a shorter delay time before droplet formation, and increased nucleation rate (slope) compared to non-seeded nucleation. **C.** Quantification of the number of condensates nucleated over time in the same cell before and after JQ1 treatment demonstrates the expected changes in delay time and slope. **D.** Representative images of a 4 by 4 micron square nuclear area as light-induced BRD4^FL^-mCh-sspB Corelet condensates nucleate rapidly in untreated conditions (2 seconds, top), but are delayed in the same cell after JQ1 treatment (10 seconds, bottom). **E.** Quantification of the nucleation rate (slope), delay time, and number density for 9 cells of similar expression level before and after JQ1 treatment. **F.** Repeated activation-deactivation cycles of BRD4^ΔN^ and overlay of condensate positions in subsequent cycles shows whether nucleation occurs repeatedly in the same nuclear locations. **G.** Overlay of condensate positions in subsequent cycles of activation for BRD4^FL^ -/+ JQ1. **H.** Pearson Correlation Coefficient (PCC) quantification and difference in PCC (deltaPCC) of overlaid images of 33 cells in subsequent activation cycles. Negative controls are PCCs between images of different cells. ***p = 0.0015 by Mann-Whitney exact, two tailed t-test. **I.** Short-term (2 minutes) 200 nM JQ1 treatment disperses BRD4 condensates (JQ1), yet they form again quickly after washing out JQ1 (Recovery). **J.** Quantification of condensate dissolution during JQ1 treatment (shaded area) and recovery after wash-out. Error bars represent standard deviation of 3 biological trials of 10 cells each. **K.** Overlay of images before/during JQ1 treatment and before treatment/after JQ1 washout (left). PCC of 30 cells before/during JQ1, and before treatment/after JQ1 washout.

To further test the concept that chromatin-binding BRD4 condensates are heterogeneously nucleated, we took advantage of the reversibility of the Corelet system to perform repeated activation-deactivation cycles of Corelet condensation and compared the nucleation positions of BRD4^ΔN^ (Fig. 4F) and BRD4^FL^ -/+ JQ1 (Fig 4G) condensates in subsequent cycles. Heterogeneous nucleation is a stochastic process biased toward forming condensates at certain seed locations, and thus should promote condensate formation preferentially at a set of nuclear locations over subsequent cycles, which would result in relatively high Pearson Correlation Coefficient (PCC) of overlaid images from multiple cycles, though inherent stochasticity will prevent the PCC from approaching 1. Conversely, homogeneous nucleation results in randomly located condensates that should exhibit lower PCC across cycles, representing randomly localized nucleation. As expected, we find low correlation between subsequent condensate positions of chromatin non-interacting BRD4^ΔN^ (Fig. 4F). Moreover, upon measuring PCC of BRD4^FL^ condensate nucleation sites in the same cell before and after JQ1 treatment, we find that condensates with chromatin-binding intact (-JQ1) exhibit higher correlation between nucleation positions in subsequent activation cycles than those without chromatin binding (+JQ1) (Fig. 4G, H). Interestingly, even the chromatin non-interacting +JQ1 case exhibits a slightly higher PCC than a negative control (measured by correlation with another cell), perhaps indicating anti-correlation with chromatin density as has been previously reported^31,37^ but this effect is much smaller than the difference between -/+JQ1 experiments. These data suggest that chromatin binding sites can act as nucleators of BRD4^FL^ condensation in living cells.

To ensure that these nucleation behaviors reflect chromatin binding and are not an artifact of Core particle binding, we investigated whether BRD4^FL^-miRFP condensates without the Corelet system form again at their initial positions after dispersal through JQ1 treatment (Fig. 4I). We observed the nuclear position of BRD4^FL^-miRFP condensates before JQ1 treatment (pre-treat), added JQ1 for 2 minutes, then washed out the JQ1 by refreshing the media twice in quick succession, and observed the dynamics and positioning of condensate reformation (Fig. 4I-K). Strikingly, condensates nucleate after JQ1 washout within seconds (Fig. 4J) and have a strong correlation with their previous locations (Fig. 4K), indicating that chromatin binding can heterogeneously seed BRD4 condensate nucleation without additional synthetic oligomerization through the Corelet system. Together these results strongly suggest BRD4^FL^ condensates have preferential nucleation sites that are likely acetylated chromatin regions.

### Coarse-grained simulations characterize BRD4 chromatin-seeded nucleation behavior

Utilizing the coarse-grained simulation system introduced in Figure 3, we can explore the molecular scale parameters that underlie the experimentally observed change in BRD4 nucleation behavior with and without chromatin-binding. A typical nucleation event in a constant-pressure simulation is characterized by following the time-dependent evolution of the largest cluster of condensing species (Fig. 5A). Simulation time is measured with respect to the time required for a Corelet to diffuse its own diameter, *τ*_d_. Initially, small clusters form and dissolve, until a cluster larger than the critical nucleus emerges stochastically and grows to form a stable condensate, *τ*_nucl_. This cluster will then continue to grow until it is large enough to be visualized in experiments after a time *τ*_delay_.

**Figure 5:**
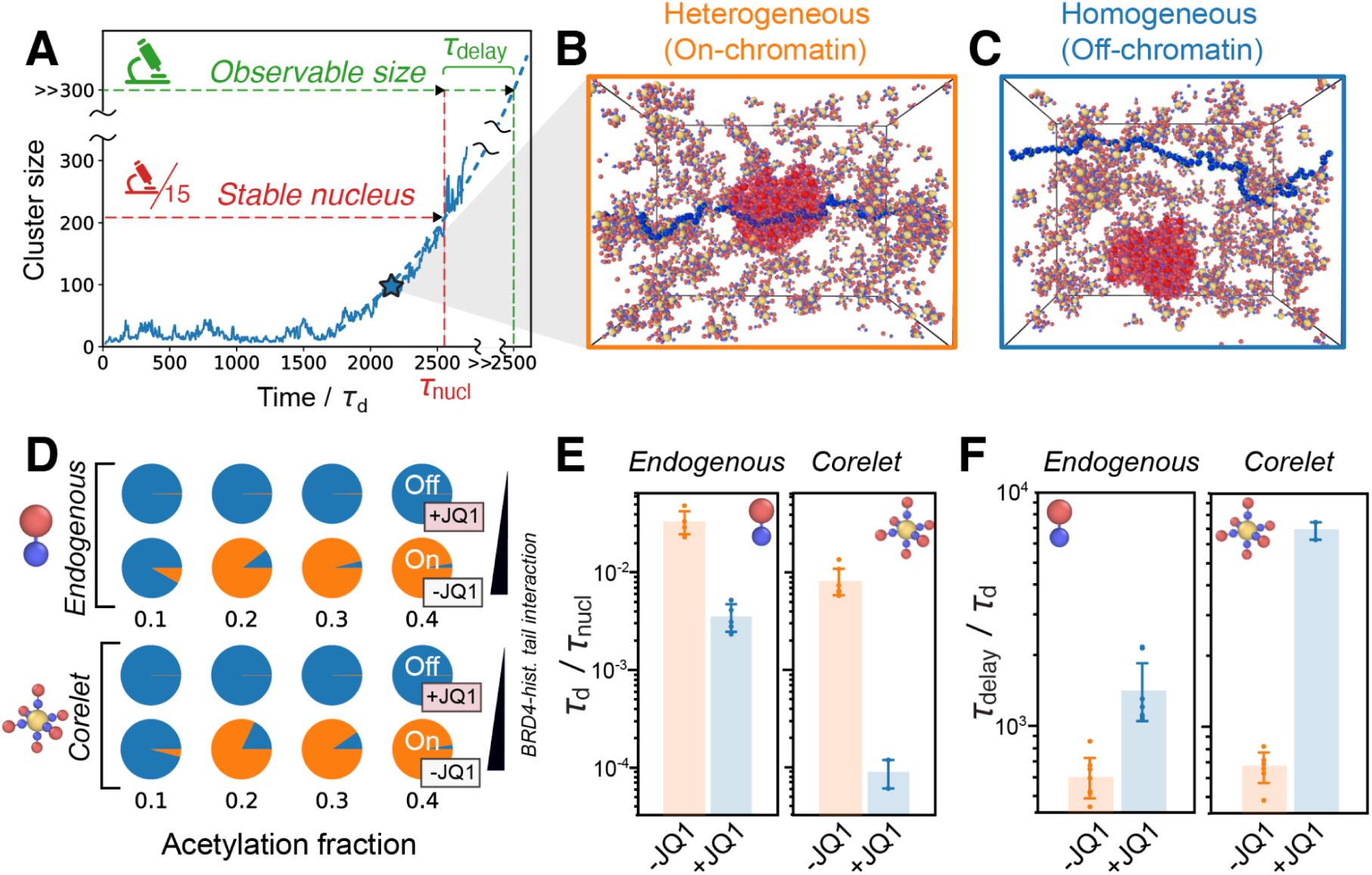
Simulations of BRD4^FL^ Corelet condensation quantify enhancement of chromatin-seeded nucleation. **A.** Schematic of a typical nucleation event. *τ*_nucl_ is the time required for cluster size to reach the threshold to form a stable nucleus (red line). *τ*_delay_ is the time required for the nucleated cluster to grow via diffusion-limited growth to reach a size that is observable under the microscope (green line), which is estimated to be 15 times larger than the stable nucleus size in our simulations. **B.** Example of a heterogeneous (on-chromatin) nucleation event, in which the largest cluster (highlighted in red) interacts with the chromatin polymer. **C.** Example of a homogeneous (off-chromatin) nucleation event. **D.** Probability of on-chromatin (orange) or off-chromatin (blue) nucleation events for endogenous (top) and Corelet (bottom) simulations at two BRD4-histone tail interaction strengths (representing +JQ1 and -JQ1), and four acetylation fractions (0.1, 0.2, 0.3, 0.4). **E.** Quantification of the nucleation rate in endogenous and Corelet simulations with (-JQ1) and without (+JQ1) strong BRD4-histone tail interactions. **F.** Quantification of the delay time before observable condensates are formed in endogenous and Corelet simulation systems with (-JQ1) and without (+JQ1) strong BRD4-histone tail interactions. In **E** and **F**, 7 independent simulations for each measurement are run at an acetylation fraction of 0.4 for both +JQ1 and -JQ1. In the +JQ1 Corelet simulations, only two out of seven simulations achieved nucleation. Error bars represent the standard error.

We expected that BRD4 condensate nucleation should depend on 1) supersaturation, 2) chromatin binding capability, and 3) the fraction of chromatin tails that are in an interaction-competent state. To this end, simulations were performed with a variable fraction of chromatin tails in the interaction-competent state, and with the condensing species chosen to be either the BRD4^FL^ Corelet (valence = 6) or the BRD4^FL^ endogenous monomer, at a series of supersaturation degrees, *S*. This BRD4^FL^ Corelet valence of 6 is above the critical valence required for off-chromatin nucleation, such that both heterogeneous nucleation (on-chromatin, Fig. 5B) and homogeneous nucleation (off-chromatin, Fig. 5C) are possible, depending on the simulation conditions.

In simulations performed with exceedingly weak attractive forces between the interaction-competent tails and BRD4, paralleling JQ1-treated cells, off-chromain nucleation occurs in 100% of simulations regardless of the fraction of interaction-competent tails (Fig. 5D, low chromatin interaction). However, if the simulations are performed with strong interactions between BRD4 and histone tails, paralleling control (-JQ1) conditions, chromatin-seeded nucleation occurs with higher probability at higher acetylation levels (Fig. 5D, high chromatin interaction strength). Remarkably, with chromatin interaction, the transition from off- to on-chromatin nucleation is a sharp function of the fraction of acetylated histone tails, and occurs between 10% and 20% of tails acetylated (Fig 5D). This implies that, on average, only one acetylated histone tail per nucleosome is necessary to shift to a majority of condensates nucleating on-chromatin in both the Corelet and endogenous BRD4 models.

We expected from classical nucleation theory that the ratio of off- to on-chromatin nucleation would also depend on the degree of supersaturation *S*, with increased *S* leading to more off-chromatin nucleation. To confirm this expectation, we repeated our simulations of the Endogenous BRD4 and Corelet systems at three degrees of supersaturation (Figure S3A). As predicted, higher supersaturation leads to more off-chromatin nucleation, and this effect is stronger with the Corelet system, due to the extra oligomerization from the Core platform.

To compare our simulations with the experimentally accessible metrics of nucleation in Figure 4, we estimated the nucleation rate, ⟨1/*τ*_nucl_⟩, and average delay time, ⟨*τ*_delay_⟩, for both models under highly acetylated conditions with either weak BRD4-chromatin interactions (+JQ1) or strong BRD4-chromatin interactions (-JQ1) (Fig. 5E-F). We find that, similar to experiments, the delay time is increased and the nucleation rate is decreased without chromatin binding for both Corelets and endogenous BRD4 model systems. Supersaturation also affects *τ*_nucl_ and *τ*_delay_, with higher supersaturation leading to lower nucleation rates and longer delay times (Fig. S3B, C). Together, these simulations state that acetylated chromatin is a potent nucleator of BRD4 condensation, that the rate of nucleation depends on the fraction of acetylated tails, and that the influence of chromatin interactions on the nucleation dynamics are highly similar for endogenous BRD4 condensation and multivalent Corelet-driven BRD4 condensation.

**Supplemental Figure 3.**
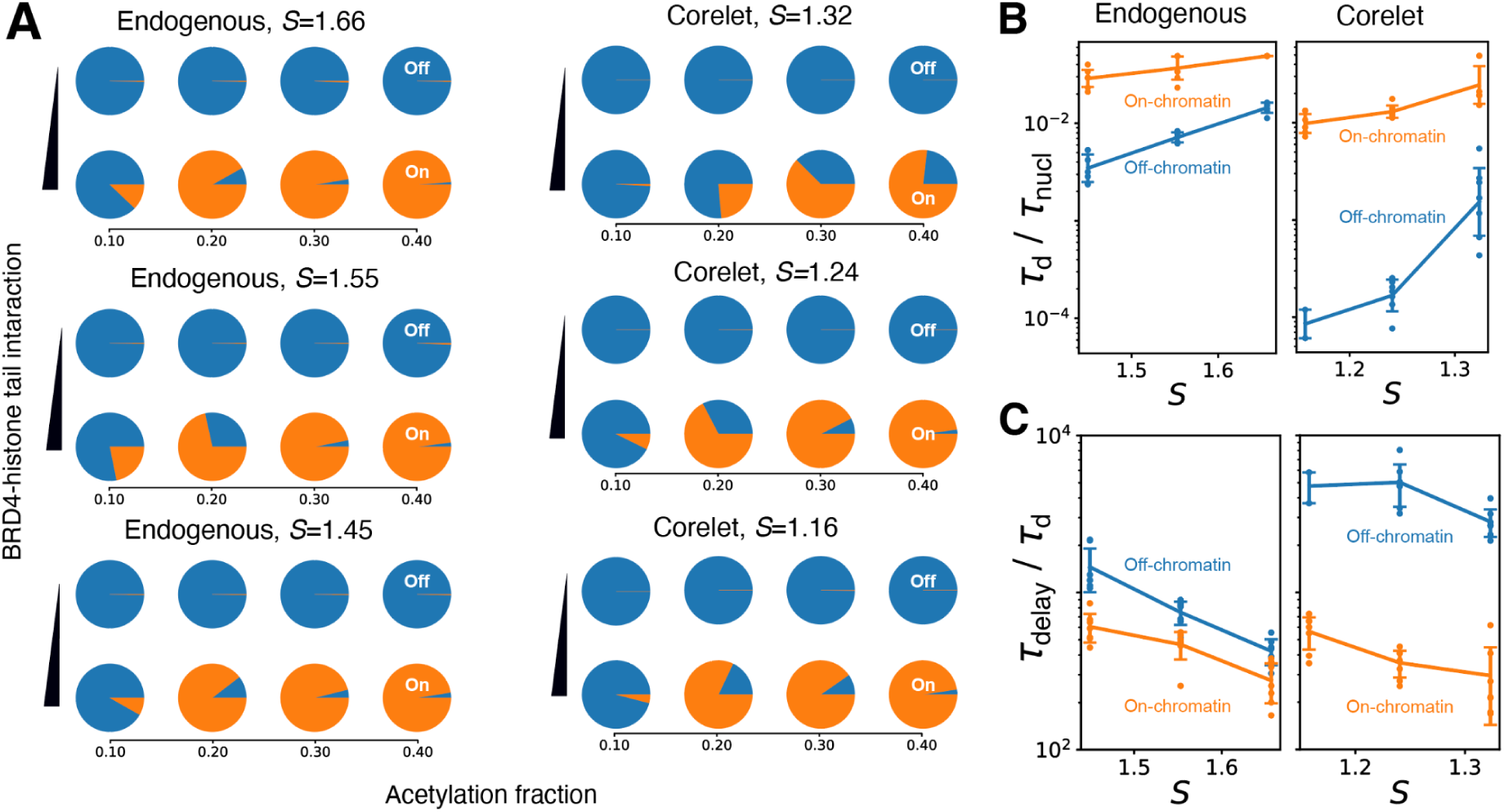
The ratio of homogeneous to heterogeneous nucleation is dependent on supersaturation, related to Figure 5. **A.** Pie charts representing the probability of on- (orange) and off-chromatin (blue) nucleation in Endogenous and Corelet simulation systems at three supersaturation levels (highest at top). Supersaturation, *S*, is defined as the ratio of the applied pressure to the equilibrium coexistence pressure. **B.** Nucleation rates and **C.** delay times as a function of the supersaturation. Error bars represent the standard error.

### Chromatin acetylation controls localized transcription factor condensation

Given that both our coarse-grained simulations and experiments suggest that acetylated regions can act as preferential nucleation sites, altering the amount of acetylated chromatin should affect the nucleation behavior of BRD4 condensates. In particular, our simulations predict that reducing acetylation levels below one acetyl per nucleosome should increase the probability of nucleating off-chromatin via the slower homogeneous pathway (Fig. 5D). To manipulate the acetylation in living cells, we treated cells with two drug treatment paradigms: 1 μM A485 for 24 hours, which inhibits p300 histone acetyltransferase and decreases the H3K27 Acetylation level to half that of WT levels, or 100 nM trichostatin A (TSA) for 24 hours, which inhibits histone deacetylases thereby increasing H3K27 Acetylation level by 3.2-fold (quantified by immunofluorescence in Fig. S4A-B).

Immunofluorescence of endogenous BRD4 at high resolution shows nuclear BRD4 colocalizing with regions of H3K27Ac (Fig. 6A, control). Upon A485 treatment, the BRD4 puncta appear larger and less colocalized with H3K27Ac, potentially representing an increase in off-chromatin nucleation. Increased amounts of H3K27Ac are clearly visible in TSA-treated cells and exhibit high colocalization with BRD4. The number of puncta per nucleus in this endogenous case represents the number density after nucleation. Quantification shows that the BRD4 puncta in A485-treated cells with lowered acetylation are less abundant (Fig. 6B) and larger (Fig. 6C) than control cells, while the BRD4 puncta in TSA-treated cells that have increased acetylation are more abundant and smaller than control, even though the endogenous expression level of BRD4 remains unchanged across these conditions (Fig. S4C). This observation is consistent with a shift in the ratio of on-chromatin and off-chromatin nucleation toward off-chromatin in the lowered acetylation case, and toward on-chromatin in the increased acetylation case.

**Figure 6.**
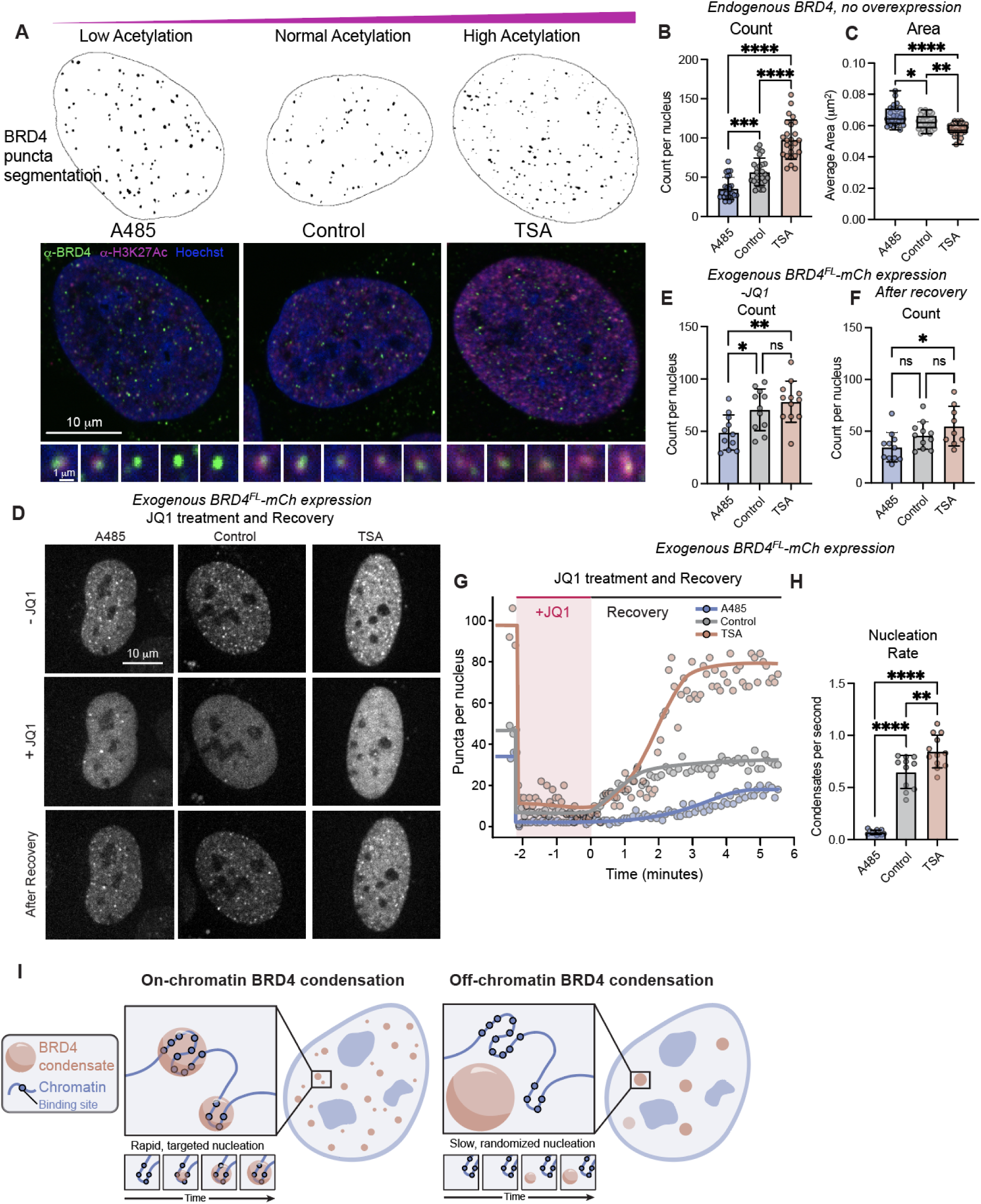
Epigenetic acetylation level influences nucleation behaviors in endogenous and exogenous BRD4 systems. **A.** Representative images and analysis examples of endogenous BRD4 puncta by immunofluorescence in cells treated with control media (Control), Histone Acetyltransferase inhibitor (A485), or histone deacetylase inhibitor (TSA). **B-C.** Quantification of endogenous BRD4 condensate count per nucleus (B) and condensate average area (C) measured by immunofluorescence in drug-treated cells. n = 25 cells each, error is SD. ***p=0.002, ****p=0.0001 by one-way ANOVA. **D.** Representative images of BRD4^FL^-mCh puncta in living cells treated with control media, A485, or TSA at three timepoints: before addition of JQ1 (-JQ1), during JQ1 incubation (+JQ1) and after JQ1 washout and recovery. **E-F.** Quantification of the number of condensates per nucleus in drug treated cells before addition of JQ1 (E) and after washout and recovery (F). Expression range as defined in Fig. S4D, E. n = 11, 11, 12 cells, error is SD, *p=0.031, **p=0.008 by one-way ANOVA. **G.** Timecourse graph of the number of BRD4^FL^-mCh puncta per nucleus during JQ1 treatment (shaded area) and nucleation after washout. **H.** Nucleation rate of BRD4^FL^-mCh puncta measured as slope after JQ1 washout. N = 11, 11, 12 for A485, control, TSA respectively. **p=0.006, ****p=0.001 **I.** Model figure summarizing findings about on- and off-chromatin BRD4 nucleation.

To directly measure the nucleation rate of BRD4^FL^-mCh condensates in TSA- and A485-treated cells without extra oligomerization from Corelets, we utilized application and wash-out of JQ1 (Fig. 6D-H, similar to Fig. 4J). In agreement with the altered number of endogenous BRD4 puncta, we observe a significantly decreased number of BRD4^FL^-mCh condensates in live A485-treated cells with lowered acetylation, and an increased number of puncta in TSA-treated cells with higher acetylation (Fig. 6D-E). Addition of 1 μM JQ1 disrupts chromatin binding and results in a rapid decrease in the number of condensates in all cells (Fig. 6D, G). After 2 minutes of JQ1 treatment, the media was washed twice in quick succession to remove the JQ1 and the rate of BRD4^FL^-mCh condensate nucleation and final number density of condensates post-recovery were recorded (Fig. 6F-G). Fitting the linear portion of the slope reveals that A485-treated cells with lower acetylation nucleated BRD4 condensates significantly slower than control, and TSA-treated cells with increased acetylation nucleated significantly faster (Fig. 6H), consistent with the predictions made by our simulations. Notably, the number density difference between A485 and TSA-treated conditions in cells exogenously expressing BRD4^FL^ is less significant than the endogenous number density difference. This likely arises due overexpression increasing the supersaturation and thereby making the system inherently less dependent on chromatin as a nucleation site.

Collectively, these results demonstrate how substrate binding through epigenetic marks can promote heterogeneously seeded condensate formation, underscoring the importance of epigenetic control over chromatin-bound condensate targeting through modulation of chromatin valence. Altogether, we demonstrate that epigenetic modification of chromatin allows for rapid condensation of transcription factors at specific chromatin sites.

**Supplemental Figure S4.**
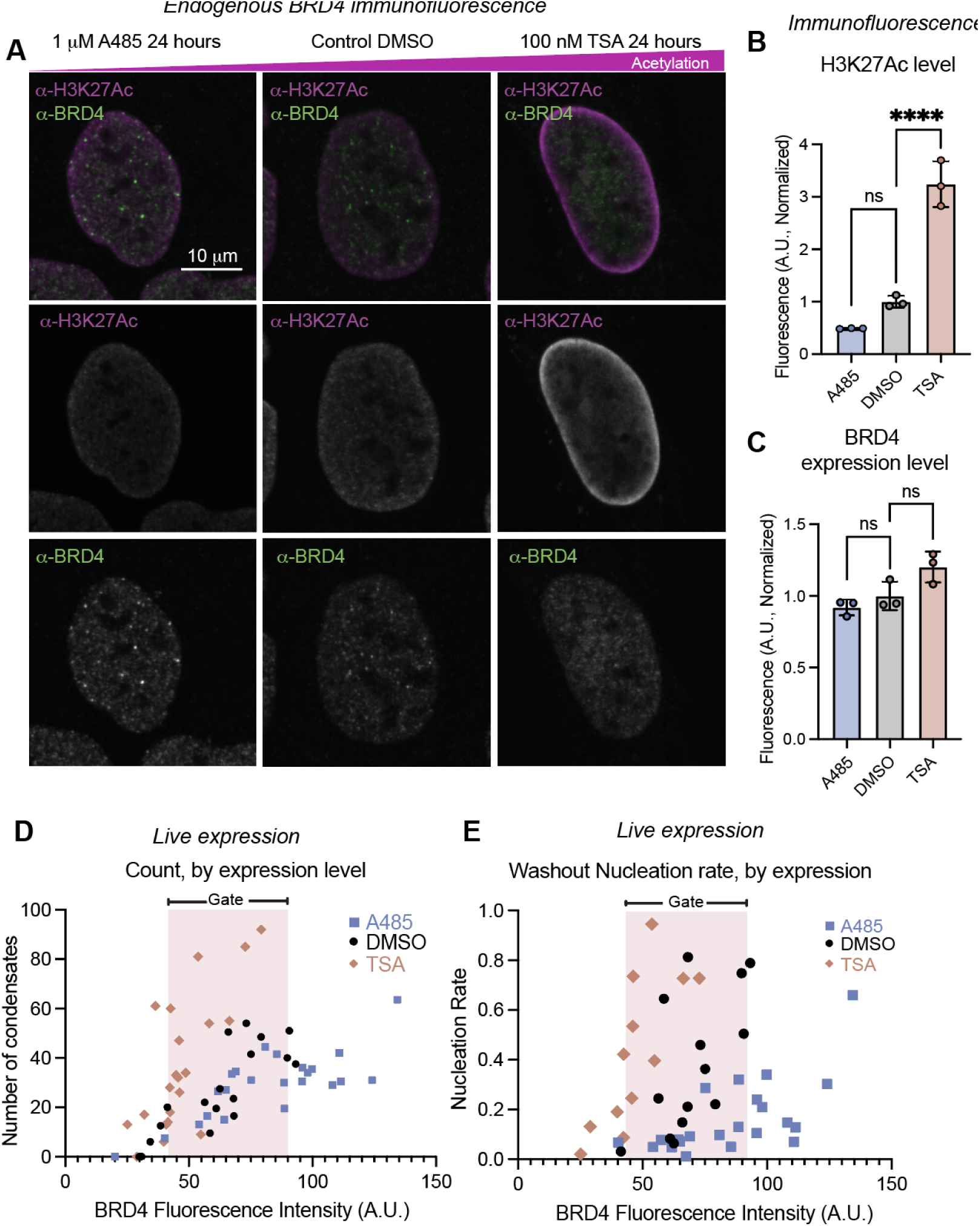
Epigenetic modifying drugs alter acetylation but not BRD4 expression level, related to Figure 6. **A.** Representative images of immunofluorescence of U2OS cells treated with DMSO (control), 1 uM A485, or 100 nM TSA for 24 hours, stained with antibodies that recognize BRD4 (green) and H3K27Ac (magenta). **B.** Quantification of H3K27 Acetylation intensity across 3 biological replicates of 25 cells each. Error is SD, ****p=0.0001 by one-way ANOVA. **C.** Quantification of BRD4 endogenous expression level across 3 biological replicates of 25 cells each. Error is SD. ns = not significant by one-way ANOVA. **D.** Count of condensates per nucleus as a function of BRD4^FL^-mCh expression level in living cells. An expression level gate is used to bound the expression level of cells for calculating average count in Fig. 6E, the same as was used in Fig. 1. **E.** Nucleation rate as a function of expression level is shown, with the expression level gate shaded.

## Discussion

Here we described the biophysical impact of chromatin binding on condensate formation by BET family transcription factor BRD4, which is biologically implicated in potent transcriptional activation, and is commonly misregulated in cancer^49–51^. We set out to investigate targeting of condensation using transcription factor BRD4 as an example of a condensation-prone protein that also binds to acetylated chromatin. We showed that BRD4’s binding to the acetylated chromatin substrate enhances both kinetic and thermodynamic properties of BRD4 condensation in three experimental systems: endogenous BRD4 distribution, exogenous BRD4-mCherry expression and synthetically oligomerized BRD4-mCherry-sspB in the Corelet system. We also developed coarse-grained simulations that represent condensation in both the Corelet and Endogenous systems, which allowed quantitative characterization of varying valence, supersaturation and acetylation fraction on condensation. With this coarse-grained model, we were able to create phase diagrams that matched the valence shift observed upon loss of chromatin binding in Corelet experiments. This also revealed that the presence of these chromatin-interacting clusters does not delineate a unique binodal boundary. Additionally, in this low-valence regime, the size of the cluster depends on the size of the acetylated region, since chromatin is a necessary, often majority, component of the cluster under these conditions. This represents a potential mechanism of controlling the size of BRD4 clusters at low expression levels, that is in agreement with experimentally observed BRD4 condensate size distribution (Fig. S2).

Even though chromatin-bound BRD4 creates non-canonical condensates with chromatin as a major component at low valence, we found that nucleation of these condensates was still consistent with the principles of classical nucleation theory through heterogeneous nucleation. Indeed, acetylated chromatin regions can act as seeds for heterogeneous nucleation of BRD4 condensates, providing rapid and targeted condensate formation at sites of epigenetic acetylation. With an overall acetylation fraction of 20%, simulated BRD4 condensates nucleate almost exclusively on-chromatin, meaning that acetylated chromatin is a potent nucleation substrate and that we expect almost all BRD4 condensates in living cells to be associated with an acetylated chromatin region. In agreement with this, repeated activation-deactivation cycles of Corelet condensation demonstrated that BRD4 condensates repeatedly form at preferred sites on chromatin, and non-Corelet BRD4-mCherry condensates nucleate again at specific locations after disruption of chromatin attachment through JQ1 treatment. Relatedly, we find that manipulation of acetylation level through epigenetic inhibitors affects the average nucleation rate of BRD4-mCherry exogenous expression condensates, as well as the number density of endogenous BRD4 droplets. These data provide a mechanism for selective formation of BRD4 condensates at acetylated regions and, excitingly, are consistent with changes to endogenous BRD4 distribution through altered nucleation pathways.

Historically, models of transcription factor targeting have been put forth that suggest mechanisms by which chromatin-binding proteins find their cognate binding sites in the large volume and complex environment of the eukaryotic nucleus^52^. These models commonly focus on describing diffusion pathways that reduce the dimensionality of the system to enable a more efficient single-molecule search for a rare binding site^53^. Here, we propose a complementary model of heterogeneously seeded transcription factor condensate nucleation (Fig. 6I), in which BRD4 condensates are rapidly heterogeneously seeded at chromatin regions enriched with acetylated binding sites, resulting in many targeted but size-limited condensates per nucleus. Disrupting chromatin binding through JQ1 treatment or overexpression of BRD4 can lead to off-chromatin nucleation, which occurs more slowly at random nuclear locations, and results in fewer, larger, and randomly localized condensates. This model explains both targeted binding of transcription factors to chromatin-localized binding sites, and depleted nucleation at off-target sites after on-target sites have nucleated. It is also consistent with published data on BRD4 targeting being dependent on its expression level^29,54–56^ and chromatin acetylation patterns^51,57,58^.

Our condensation model of BRD4 chromatin-targeting provides context for altered BRD4 chromatin localization patterns observed in disease states due to aberrant acetylation patterns^59,60^. Interestingly, the BET inhibitor JQ1 was originally developed as an anti-cancer therapy for tumors of various origins to prevent transcriptional expression of oncogenes that become hyperacetylated^36^. We found that JQ1 treatment disrupts chromatin binding, as expected, but does not completely abrogate BRD4 condensation ability–instead, JQ1 treatment leads to randomized localization of BRD4 nucleation. Similarly, other small molecule drugs that target chromatin-interaction motifs may modulate but not abrogate condensation of transcription factors, and therefore lead to variable biological effects.

The modulation of condensation behaviors through chromatin binding that we observe with BRD4 may represent a general phenomenon that applies to other BET family proteins or even other transcription factor families. We expect that transcription factors with dual chromatin binding and condensation capabilities like BRD4 will follow the kinetic and thermodynamic paradigms described here, including chromatin-bound clustering at low valence and concentration, as well as heterogeneously seeded nucleation at multivalent chromatin binding regions. Overall, our results put forth a model in which the cell utilizes epigenetic modifications to regulate condensation properties of transcription factors in order to target their nucleation and enact a specific gene expression profile.

## Supporting information

Supplemental simulation parameters

## Acknowledgements

We thank David Sanders for his generous gifts of plasmids, and members of Brangwynne and Jacobs laboratories at Princeton University for helpful feedback and discussion throughout the project. A.R.S. would like to thank Jennifer M. Leija for lively conversation during long microscopy sessions. This work was funded in part by the National Science Foundation DMR-2143670 (W.M.J), AFOSR FA9550-20-1-0241 (C.P.B.), the St. Jude Research Collaborative on the Biology and Biophysics of RNP granules (C.P.B.), and the Howard Hughes Medical Institute (C.P.B.). A.R.S. was supported by a Life Science Research Foundation Fellow through the Mark Foundation for Cancer Research (AWD1006303) and an NIH Pathway to Independence award (NCI K99 AWD1007843). J.M.E. was supported by a Rubicon Grant (NWO). C.J.W. was supported by an NSF GRFP (DGE-2039656).

## Author Contributions

A.R.S. and J.M.E. conceived of and performed most experiments, with advice from C.P.B. C.J.W performed and analyzed experiments using TSA and A485, with advice from C.P.B.. Y.P. and H.W. developed simulations with advice from W.J.. D.B., N.D.O. and C.C.J. performed preliminary experiments, quantitative microscope calibrations and associated controls vital to the initiation of the project.

## Methods

### Cell culture

U2OS cells, obtained from the ATCC and authenticated using ATCC’s STR profiling, were cultured in DMEM (GIBCO, 11995065) supplemented with 10% FBS (Atlanta Biological, S11150H) and 1% streptomycin and penicillin (GIBCO, 15140122). The cells were grown under conditions of 37°C with 5% CO_2_ and tested for mycoplasma regularly.

### Immunofluorescence

Cells plated in 96 well plates were fixed by adding 4% paraformaldehyde for 10 minutes. Following fixation, cells were subjected to a 5-minute wash with buffer (0.35% Triton-X, Thermo Fisher PRH5142, in PBS) and permeabilized with 0.5% Triton-X in PBS for 1 hour. Subsequently, cells were blocked for 1 hour using blocking buffer (0.25% Triton-X, 5% FBS, in PBS). Primary antibodies (anti-BRD4, Cell Signaling Technologies Mouse mAb #63759 at 1:1000 dilution, anti-H3K27ac, Active Motif #39133, 1:1000 dilution) were prepared in blocking buffer and incubated on the sample overnight at 4 °C. The following day, cells were washed 3 times for 5 minutes each with washing buffer. The secondary antibodies (AlexaFluor 647 goat-anti-rabbit Thermo Fisher A-21245, 1:1000 dilution) were prepared in blocking buffer and incubated for 2 hours at 4 °C. Cells were then washed 3 times for 5 minutes each with wash buffer, followed by a 20-minute incubation with Hoechst (1:2000 dilution). Finally, cells were washed with PBS.

### Construct design and cloning

The full length BRD4 DNA fragment was amplified by PCR from pcDNA4-TO-HA-BRD4FL (Addgene plasmid #31351)^61^ incorporated into linearized FM5 lentiviral vectors containing standardized linkers (generously provided by David Sanders) using the In-Fusion HD cloning kit (Takara Bio, 638910). BRD4^dN^-mCh-sspB (Addgene plasmid #121968)^31^ and NLS-iLID-Ferritin (Addgene plasmid #122147)^40^ were originally developed and characterized in previous Brangwynne lab studies. The validity of all constructs was verified through GENEWIZ (Azenta Life Sciences) plasmid Sanger sequencing.

### Lentivirus production and lentiviral transduction

For all live cell experiments, cells were stably transduced with lentivirus. Lentiviruses were generated by seeding Lenti-X 293T cells (Takara Bio, Cat. No. 632180) in 6-well plates, at approximately 70% confluence at the time of transfection. After 24-48 hours, the transfer plasmid and helper plasmids VSV-G and PSP (generously provided by David Sanders) were transfected into the Lenti-X cells using Transit293 transfection reagent (Mirus, Cat. No. MIR 2700) and incubated in OptiMEM (ThermoFisher Cat No. 31985062). Approximately 48 hours after transfection, the viruses were harvested and filtered using a 0.45 μm filter (VWR Cat. No 28144-007). The resulting viruses were either used immediately or stored at -80°C. U2OS cells were plated in 96-well glass-bottom plates (ThermoFisher Cat. No NC0536760) at 30-50% confluency and transduced with lentivirus for 2-3 days prior to live-cell imaging experiments.

### Drug treatments

JQ1 (MedChemExpress, Cat. No. HY-13030) diluted in DMSO was added to the cells to a final concentration of 1 μM, and incubated for 90 minutes prior to imaging. The +JQ1 cells were imaged in media containing JQ1. Trichostatin A (TSA, Sigma, T8552-1MG) was resuspended in DMSO and added to the cells to a final concentration of 100 nM, and incubated for 24 hours prior to imaging. A485 (MedChemExpress, Cat. No. HY-107455) was resuspended in DMSO and added to the cells to a final concentration of 200 nM, and incubated for 24 hours prior to imaging.

### Corelet system optogenetic activation

Cells expressing Corelet system constructs (NLS-iLID-GFP-Ferritin and either BRD4dN-mCh-sspB or BRD4FL-mCh-sspB) via lentiviral transduction were seeded on 96-well glass bottom plates (ThermoFisher Cat. No NC0536760), then loaded onto a Nikon A1 laser scanning confocal microscope with Nikon Eclipse Ti2 body. A 100X oil immersion Apo TIRF objective (NA 1.49) was used to visualize these samples. Obtaining images with the 488 nm laser at 0.1% laser power and above is sufficient to trigger iLID-sspB interaction and Corelet system optogenetic activation. Activation movies were obtained with 488 nm and 512 nm lasers every 5 seconds for 3 minutes.

### Image analysis for count, size, total area, volume of puncta per nucleus

Image analysis was performed in FIJI^62^, an open-source image analysis platform. Nuclei were initially segmented from single z-plane images of BRD4^FL^-mCh alone (Fig. 1) or from the final frame of Corelet Activation movies (Fig. 2) using the mCherry (561 nm laser) channel and an Otsu thresholding method, with smoothing if necessary to obtain a nuclear mask. Puncta were segmented within each nucleus using theIsoData method in FIJI after smoothing and rolling ball background subtraction with a radius of 4 pixels. The Analyze Particles feature was used to measure the area, count and intensity of pixels within puncta for each nucleus. Graphs of puncta Count were created by averaging the number of puncta per nucleus across 25 nuclei within a well (biological replicate), then across four biological replicates. From the same data, puncta per nucleus were also plotted on a cell-by-cell basis on a log-log plot before and after addition of JQ1 in the same nucleus (Fig. S2D). Graphs of puncta Average Size were created by first averaging the average area of puncta within a nucleus, then across 25 nuclei within a well (cells within one well consisting of a biological replicate), then across four biological replicates. Condensate Volume was estimated by raising the average area of condensate to the power of 3/2, then multiplying this by the number of condensates per nucleus.

### Phase Diagram data acquisition

The Nikon A1 laser scanning confocal microscope is equipped with PicoQuant software and was calibrated by FCS at a range of 488 nm and 561 nm laser settings from 0.1-5% power at 512×512 or 1024×1024 pixels per image to create a quantitative conversion from fluorescence arbitrary units (AU) of GFP and mCherry fluorophores to micromolar concentrations of these proteins. U2OS cells expressing Corelet system components (NLS-GFP-iLID-Ferritin and BRD4FL-mCh-sspB) were loaded onto the Nikon A1 laser scanning microscope. To ensure consistency, cells were allowed to equilibrate to stage temperature for 30 minutes before imaging, and a standardized imaging protocol was applied to all cells used in the phase diagram. Activation movies were obtained every 2 seconds for 3 minutes with 488 nm and 512 nm lasers. Positions of acquired cells were marked, then media containing JQ1 to a final concentration of 1 uM was added to the imaged well and incubated for 90 minutes on the stage-top. A second round of activation series of images every 2 seconds for 3 minutes were taken of all cells, used for +JQ1 data.

### Corelet system image analysis and phase diagram construction

Only nuclei fully within the field of view were included in the analysis. Nuclei were segmented using ImageJ Otsu method, and the average fluorescence intensity of GFP and mCherry was measured based on the first frame of the movie, before activation. The intensity measurement in AU was converted to micromolar concentration via the calibrated imaging settings. Cells were marked as ‘Yes PS’ or ‘No PS’ through qualitative assessment of the first and last frames of the 3 minute activation series, determined by whether new condensates were formed during the activation period. Qualitative assessments were conducted by two independent observers, with one observer blinded to the experimental conditions. The evaluations of the two observers showed high consistency in almost all cases. Instances where the observers disagreed on cell classification were excluded from the results. Core concentration for each nucleus was calculated as the GFP concentration divided by 24 (as there are 24 GFP monomers for each assembled Ferritin Core). Valence of each nucleus was calculated as the ratio of sspB-fused protein (mCherry concentration) to Core (GFP concentration of 24-mers). Data were plotted in GraphPad Prism 10, and shaded area was added manually in Adobe Illustrator to guide the eye.

### Nucleation measurements

U2OS cells expressing Corelet system components (NLS-GFP-iLID-Ferritin and BRD4FL-mCh-sspB) were imaged on a Nikon X1 spinning disc confocal (described above) in 488 and 561 channels every 2 seconds for 5 minutes. The number of condensates per nucleus were counted in each frame through image analysis in FIJI, with smoothing and rolling ball background subtraction radius 4, then thresholded with IsoData algorithm and counted per nucleus using the Analyze Particles feature.

#### Slope, delay time and number density

Nucleation Slope was determined by fitting the rate of puncta per nucleus formed over the linear portion of the curve. For -JQ1 samples, the linear portion of the curve usually consisted of the first 3-4 frames and for +JQ1 samples, the linear portion was generally shifted later. Delay time was determined as the time in seconds before the number of puncta increased consistently for 2 frames. Number density was determined as the count of puncta per nucleus at the last frame of the activation series (5 minutes of activation).

### Repeated cycles of Corelet activation and deactivation

U2OS cells expressing Corelet system components (NLS-GFP-iLID-Ferritin and BRD4FL-mCh-sspB) were imaged on a Nikon X1 spinning disc confocal (described above) for five segments: Activation-Deactivation-Activation-Deactivation-Activation. “Activation” with 488 and 561 channels imaged every 2 seconds for 3 minutes, “Deactivation” with only 561 channel imaged every 2 seconds for 5 minutes (absence of 488 nm laser allows deactivation of iLID-sspB interaction).

#### Pearson Correlation Coefficient

Activation-deactivation cycling series were registered in FIJI using HyperStackReg, Rigid Body, to account for cell movement. The last frame of each of 3 activation cycles of the registered series was extracted, and Pearson Correlation Coefficient was run between 4×4 micron square areas of the same cell (experiment) and different cells (negative control) using the JaCoP plugin in FIJI. Correlation between cycles 1&2 and cycles 2&3 are both reported.

### JQ1 washout experiments

U2OS cells were lentivirally transduced with BRD4FL-mCh-sspB, plated on 96-well glass bottom plates and imaged 48-72 hours after transduction. Plates were loaded onto a Nikon spinning disc microscope (Yokogawa CSU-X1) on a Nikon Eclipse Ti body and DU-897 EMCCD camera and LU-NV laser launch, with stage top incubation of 37 °C and 5% CO_2_ conditions maintained by Okolab microscope stage incubator with 96-well insert. A 100X oil immersion Apo TIRF objective (NA 1.49 MRD01991) was used to obtain images with 488 nm and 561 nm lasers to visualize GFP and mCherry constructs, respectively. Movies were obtained once in each well at a frame rate of one image every 5 seconds for 10 min, with the first three frames (15 seconds) as ‘pre-treatment’ to establish the initial number of condensates per nucleus. After the first three frames, media containing JQ1 was added to the open top of the well to a final concentration of 100 nM JQ1 and cells imaged for 24 frames (2 minutes), called ‘JQ1 treatment’. After 2 minutes of JQ1 treatment, the media was removed and replaced with fresh media (no drug) twice in quick succession, then imaged until a total of 10 minutes, during which time the BRD4 condensates nucleated again, called ‘recovery’. This process was repeated in each well, with cells in the same frame considered technical replicates and cells in different wells considered biological replicates.

#### Washout Pearson Correlation Coefficient

JQ1 washout series images were registered to correct for cell movement using HyperStackReg in FIJI. Frames from the end of JQ1 treatment and end of recovery were compared in the JaCoP plugin for ‘during JQ1’ PCC, and frames from the pre-treatment and end of recovery period were compared for ‘Recovery’ PCC.

### Simulations

#### Coarse-grained model

The model for a BRD4 protein consists of 2 blobs representing the globular domain-containing (N-terminal) and disordered (C-terminal) portions of the protein. Corelets are constructed by attaching BRD4 molecules to a particle representing the ferritin core. Differential valence is represented by modifying the number of BRD4 molecules attached to the core particle in each simulation. Chromatin is represented by a chain of nucleosome particles, each of which is decorated with 8 small blobs representing the histone tails. Histone tails were each assigned one of two states: a non-acetylated state, in which they do not interact with BRD4, or an acetylated state, in which they are highly attractive to the N-terminal blob of a BRD4 molecule.

There are 3 main types of interactions in the system:

1. Steric repulsion is represented by a WCA repulsive pair potential^63^. For the BRD4 N-terminal, C-terminal, and histone tail blobs, the WCA diameters are chosen to enforce a maximum packing fraction consistent with the total volume occupied by the amino acids comprising each of the blobs. For the ferritin core and histone particles, the WCA diameters are chosen based on structural data.
2. Bonds between nucleosomes, between the N- and C-terminal polymer blobs of BRD4, and between BRD4 molecules and the oligomerization platform cores are represented using FENE potentials^64^. Spring constants and equilibrium bond lengths for bonds between polymer blobs are chosen based on an ideal polymer model.
3. Interactions between polymer blobs are modeled using the Flory-Krigbaum potential^65–67^, which describes polymers in dilute solution in terms of their radii of gyration and a variable strength of the interaction. Radii of gyration are estimated based on an ideal polymer model, and interaction strengths are chosen to reproduce the experimental phase diagram.

All MD simulations were run using the LAMMPS package^68^.

#### Phase diagram calculations

The measurement of coexistence densities and the determination of phase separation in the presence or absence of chromatin were accomplished through direct coexistence simulations. All simulations were performed in the NVT ensemble with the Nosé–Hoover thermostat and used a 120 ✕ 120 ✕ 561 nm^3^ orthogonal box with periodic boundary conditions in all 3 directions. For simulations without chromatin, 343 BRD4^FL^ Corelets were set up to be in slab geometry, which is a liquid (dense) phase slab in the x-y plane, so as to create two flat interfaces. For simulations with chromatin present, slab systems with 343, 86, and 43 Corelets were created to test multiple Corelet densities. Chromatin was constructed as a 400-nucleosome chain along the z-axis, passing through the liquid (dense) phase slab of Corelets. The opposite ends of the chromatin are bonded to each other through the z-axis periodic boundary. Measurements of coexistence densities in the absence of chromatin were obtained by analyzing the distribution of Corelets across the z-axis, from which the density in the dilute and dense regions can be determined. To identify phase separation in the presence of chromatin, we examined the distribution of Corelets along the z-axis of the simulation box. Steady-state clustering, resulting in regions of high Corelet concentration that do not dissolve over the course of the simulation, was taken as evidence of equilibrium condensate formation. Condensate formation was also verified via qualitative assessment of snapshots taken throughout the simulation.

#### Nucleation simulations

Nucleation simulations were conducted with periodic boundary conditions, where the z-direction is percolated by periodic chromatin. The x and y-directions were held at constant pressure to enforce approximately constant supersaturation in the process of condensation. The simulation box dimension in the z-direction was kept fixed to avoid unphysical coupling of the chromatin tension to fluctuations in the other box dimensions. Langevin dynamics were then used to account for an implicit solvent. Nucleation simulations were conducted using either 512 Corelets or 5832 endogenous BRD4 molecules. The number of simulated nucleosomes varies in the range ∼330-360 depending on the supersaturation. Nucleation was monitored by clustering all molecules based on a cutoff distance, *R*_clust_, and tracking the number of molecules in the biggest cluster, *N*_CS_. The cutoff distances of *R*_clust_ = 25 nm for Corelets and 12.5 nm for endogenous BRD4 were chosen based on the distance at which the nonbonded interactions between pairs of molecules are negligible for any molecular orientation.

Nucleation transition paths are defined as the first-passage portion of a trajectory in which the largest cluster in the system grows to become a stable droplet. Specifically, a transition path begins when *N*_CS_ increases beyond fluctuations around the initial metastable state and passes directly to a stable cluster size of ∼200 Corelets or ∼600 BRD4 molecules. These stable cluster sizes correspond to approximately 1/15 and 1/75 of the smallest registerable droplet sizes in the experiments, as determined from the experimental pixel dimensions and the condensed-phase densities obtained from equilibrium simulations. The first passage time, *τ*_nucl_, is recorded as the time required to reach the stable droplet size, and the nucleation rate is defined as *J* = ⟨1 / *τ*_nucl_⟩. The delay time *τ*_delay_ is obtained from the intersection of the extrapolated transition path, assuming a diffusion-limited growth law 3/2 𝑁 (𝑡) = 𝑁 (1 + 𝑏𝑡)^69^, and the smallest registerable droplet size in the experiments.

On-chromatin and off-chromatin nucleation and growth regimes were distinguished based on the average chromatin coverage by the largest Corelet or BRD4 cluster in the system, which is computed based on a cutoff distance of *R*_cut_ = *R*_clust_ = 25 nm between nucleosomes and Corelet clusters or 12.5 nm between nucleosomes and BRD4 clusters, respectively.

## Code availability

All simulations were performed using the LAMMPS molecular dynamics package, which is publicly accessible.

## References

1. Saini, B. & Mukherjee, T. K. Biomolecular Condensates Regulate Enzymatic Activity under a Crowded Milieu: Synchronization of Liquid–Liquid Phase Separation and Enzymatic Transformation. J. Phys. Chem. B (2023) doi:10.1021/acs.jpcb.2c07684.

2. Enzymatic Reactions inside Biological Condensates. J. Mol. Biol. 433, 166624 (2021).

3. Falahati, H., Pelham-Webb, B., Blythe, S. & Wieschaus, E. Nucleation by rRNA Dictates the Precision of Nucleolus Assembly. Curr. Biol. 26, 277–285 (2016).

4. Falahati, H. & Wieschaus, E. Independent active and thermodynamic processes govern the nucleolus assembly in vivo. Proc. Natl. Acad. Sci. U. S. A. 114, 1335–1340 (2017).

5. Xing, Y., Johnson, C. V., Moen, P. T., Jr, McNeil, J. A. & Lawrence, J. Nonrandom gene organization: structural arrangements of specific pre-mRNA transcription and splicing with SC-35 domains. J. Cell Biol. 131, 1635–1647 (1995).

6. Xing, Y., Johnson, C. V., Dobner, P. R. & Lawrence, J. B. Higher level organization of individual gene transcription and RNA splicing. Science vol. 259 1326–1330 (1993).

7. Moen, P. T., Jr et al. Repositioning of muscle-specific genes relative to the periphery of SC-35 domains during skeletal myogenesis. Mol. Biol. Cell 15, 197–206 (2004).

8. Brown, J. M. et al. Association between active genes occurs at nuclear speckles and is modulated by chromatin environment. J. Cell Biol. 182, 1083–1097 (2008).

9. Patil, A. et al. A disordered region controls cBAF activity via condensation and partner recruitment. Cell 186, 4936–4955.e26 (2023).

10. Zhang, Y. et al. Nuclear condensates of p300 formed though the structured catalytic core can act as a storage pool of p300 with reduced HAT activity. Nat. Commun. 12, 4618 (2021).

11. Ma, L. et al. Co-condensation between transcription factor and coactivator p300 modulates transcriptional bursting kinetics. Mol. Cell 81, 1682–1697.e7 (2021).

12. Sabari, B. R. et al. Coactivator condensation at super-enhancers links phase separation and gene control. Science 361, (2018).

13. Boija, A. et al. Transcription Factors Activate Genes through the Phase-Separation Capacity of Their Activation Domains. Cell 175, 1842–1855.e16 (2018).

14. Lyons, H. et al. Functional partitioning of transcriptional regulators by patterned charge blocks. Cell 186, 327–345.e28 (2023).

15. Villegas, J. A., Heidenreich, M. & Levy, E. D. Molecular and environmental determinants of biomolecular condensate formation. Nat. Chem. Biol. 18, 1319–1329 (2022).

16. Banani, S. F., Lee, H. O., Hyman, A. A. & Rosen, M. K. Biomolecular condensates: organizers of cellular biochemistry. Nat. Rev. Mol. Cell Biol. 18, 285–298 (2017).

17. Farag, M., Borcherds, W. M., Bremer, A., Mittag, T. & Pappu, R. V. Phase separation of protein mixtures is driven by the interplay of homotypic and heterotypic interactions. Nat. Commun. 14, 5527 (2023).

18. Shin, Y. Rich Phase Separation Behavior of Biomolecules. Mol. Cells 45, 6–15 (2022).

19. Kashchiev, D. Nucleation: Basic Theory with Applications. (Butterworth Heinemann, 2000).

20. Shimobayashi, S. F., Ronceray, P., Sanders, D. W., Haataja, M. P. & Brangwynne, C. P. Nucleation landscape of biomolecular condensates. Nature 599, 503–506 (2021).

21. Wei, M.-T. et al. Nucleated transcriptional condensates amplify gene expression. Nat. Cell Biol. 22, 1187–1196 (2020).

22. Wiegand, T. & Hyman, A. A. Drops and fibers - how biomolecular condensates and cytoskeletal filaments influence each other. Emerg Top Life Sci 4, 247–261 (2020).

23. Chayen, N. E., Saridakis, E. & Sear, R. P. Experiment and theory for heterogeneous nucleation of protein crystals in a porous medium. Proc. Natl. Acad. Sci. U. S. A. 103, 597–601 (2006).

24. Liquid Nuclear Condensates Mechanically Sense and Restructure the Genome. Cell 176, 1518 (2019).

25. Plys, A. J. & Kingston, R. E. Dynamic condensates activate transcription. Science vol. 361 329–330 (2018).

26. Wagh, K., Garcia, D. A. & Upadhyaya, A. Phase separation in transcription factor dynamics and chromatin organization. Curr. Opin. Struct. Biol. 71, 148–155 (2021).

27. Dey, A., Chitsaz, F., Abbasi, A., Misteli, T. & Ozato, K. The double bromodomain protein Brd4 binds to acetylated chromatin during interphase and mitosis. Proc. Natl. Acad. Sci. U. S. A. 100, 8758–8763 (2003).

28. Itzen, F., Greifenberg, A. K., Bösken, C. A. & Geyer, M. Brd4 activates P-TEFb for RNA polymerase II CTD phosphorylation. Nucleic Acids Res. 42, 7577–7590 (2014).

29. Wang, N., Wu, R., Tang, D. & Kang, R. The BET family in immunity and disease. Signal Transduct Target Ther 6, 23 (2021).

30. Han, X. et al. Roles of the BRD4 short isoform in phase separation and active gene transcription. Nat. Struct. Mol. Biol. 27, 333–341 (2020).

31. Shin, Y. et al. Liquid Nuclear Condensates Mechanically Sense and Restructure the Genome. Cell 175, 1481–1491.e13 (2018).

32. Wang, C. et al. The C-terminal low-complexity domain involved in liquid-liquid phase separation is required for BRD4 function in vivo. J. Mol. Cell Biol. 11, 807–809 (2019).

33. Morin, J. A. et al. Sequence-dependent surface condensation of a pioneer transcription factor on DNA. Nat. Phys. 18, 271–276 (2022).

34. Maharana, S. et al. RNA buffers the phase separation behavior of prion-like RNA binding proteins. Science 360, 918–921 (2018).

35. Niaki, A. G. et al. Loss of Dynamic RNA Interaction and Aberrant Phase Separation Induced by Two Distinct Types of ALS/FTD-Linked FUS Mutations. Mol. Cell 77, 82–94.e4 (2020).

36. Filippakopoulos, P. et al. Selective inhibition of BET bromodomains. Nature 468, 1067–1073 (2010).

37. Lee, D. S. W. et al. Size distributions of intracellular condensates reflect competition between coalescence and nucleation. Nat. Phys. 19, 586–596 (2023).

38. Feric, M. & Brangwynne, C. P. A nuclear F-actin scaffold stabilizes ribonucleoprotein droplets against gravity in large cells. Nat. Cell Biol. 15, 1253–1259 (2013).

39. Caragine, C. M., Haley, S. C. & Zidovska, A. Surface Fluctuations and Coalescence of Nucleolar Droplets in the Human Cell Nucleus. Phys. Rev. Lett. 121, 148101 (2018).

40. Bracha, D. et al. Mapping Local and Global Liquid Phase Behavior in Living Cells Using Photo-Oligomerizable Seeds. Cell 176, 407 (2019).

41. Sanders, D. W. et al. Competing Protein-RNA Interaction Networks Control Multiphase Intracellular Organization. Cell 181, 306–324.e28 (2020).

42. Louis, A. A., Bolhuis, P. G., Hansen, J. P. & Meijer, E. J. Can polymer coils Be modeled as ‘Soft colloids’? Phys. Rev. Lett. 85, 2522–2525 (2000).

43. Dignon, G. L., Zheng, W., Kim, Y. C., Best, R. B. & Mittal, J. Sequence determinants of protein phase behavior from a coarse-grained model. PLoS Comput. Biol. 14, e1005941 (2018).

44. Bianchi, E., Largo, J., Tartaglia, P., Zaccarelli, E. & Sciortino, F. Phase Diagram of Patchy Colloids: Towards Empty Liquids. Phys. Rev. Lett. 97, 168301 (2006).

45. Broedersz, C. P. et al. Condensation and localization of the partitioning protein ParB on the bacterial chromosome. Proc. Natl. Acad. Sci. U. S. A. 111, 8809–8814 (2014).

46. Erdel, F. & Rippe, K. Formation of Chromatin Subcompartments by Phase Separation. Biophys. J. 114, 2262–2270 (2018).

47. Hensley, A., Jacobs, W. M. & Rogers, W. B. Self-assembly of photonic crystals by controlling the nucleation and growth of DNA-coated colloids. Proc. Natl. Acad. Sci. U. S. A. 119, (2022).

48. Hensley, A., Videbæk, T. E., Seyforth, H., Jacobs, W. M. & Rogers, W. B. Macroscopic photonic single crystals via seeded growth of DNA-coated colloids. Nat. Commun. 14, 4237 (2023).

49. Mehta, S. & Zhang, J. Liquid-liquid phase separation drives cellular function and dysfunction in cancer. Nat. Rev. Cancer 22, 239–252 (2022).

50. Tang, S. C., Vijayakumar, U., Zhang, Y. & Fullwood, M. J. Super-Enhancers, Phase-Separated Condensates, and 3D Genome Organization in Cancer. Cancers 14, (2022).

51. Donati, B., Lorenzini, E. & Ciarrocchi, A. BRD4 and Cancer: going beyond transcriptional regulation. Mol. Cancer 17, 164 (2018).

52. Lu, F. & Lionnet, T. Transcription Factor Dynamics. Cold Spring Harb. Perspect. Biol. 13, (2021).

53. Jana, T., Brodsky, S. & Barkai, N. Speed-Specificity Trade-Offs in the Transcription Factors Search for Their Genomic Binding Sites. Trends Genet. 37, 421–432 (2021).

54. Liang, Y., Tian, J. & Wu, T. BRD4 in physiology and pathology: BET’’ on its partners. Bioessays 43, e2100180 (2021).

55. Drumond-Bock, A. L. et al. Increased expression of BRD4 isoforms long (BRD4-L) and short (BRD4-S) promotes chemotherapy resistance in high-grade serous ovarian carcinoma. Genes Cancer 14, 56–76 (2023).

56. Maruyama, T. et al. A Mammalian bromodomain protein, brd4, interacts with replication factor C and inhibits progression to S phase. Mol. Cell. Biol. 22, 6509–6520 (2002).

57. Alekseyenko, A. A. et al. Ectopic protein interactions within BRD4-chromatin complexes drive oncogenic megadomain formation in NUT midline carcinoma. Proc. Natl. Acad. Sci. U. S. A. 114, E4184–E4192 (2017).

58. Sudarshan, D. et al. Recurrent chromosomal translocations in sarcomas create a megacomplex that mislocalizes NuA4/TIP60 to Polycomb target loci. Genes Dev. 36, 664–683 (2022).

59. Lovén, J. et al. Selective inhibition of tumor oncogenes by disruption of super-enhancers. Cell 153, 320–334 (2013).

60. French, C. A. et al. BRD4-NUT fusion oncogene: a novel mechanism in aggressive carcinoma. Cancer Res. 63, 304–307 (2003).

61. Rahman, S. et al. The Brd4 extraterminal domain confers transcription activation independent of pTEFb by recruiting multiple proteins, including NSD3. Mol. Cell. Biol. 31, 2641–2652 (2011).

62. Schindelin, J., et al. Fiji: an open-source platform for biological-image analysis. Nat. Methods 9, 676–682 (2012).

63. Weeks, J. D., Chandler, D. & Andersen, H. C. Role of repulsive forces in determining the equilibrium structure of simple liquids. J. Chem. Phys. 54, 5237–5247 (1971).

64. Kremer, K. & Grest, G. S. Dynamics of entangled linear polymer melts: A molecular-dynamics simulation. J. Chem. Phys. 92, 5057–5086 (1990).

65. Flory, P. J. & Krigbaum, W. R. Statistical mechanics of dilute polymer solutions. II. J. Chem. Phys. 18, 1086–1094 (1950).

66. Lenart, P. J., Jusufi, A. & Panagiotopoulos, A. Z. Effective potentials for 1:1 electrolyte solutions incorporating dielectric saturation and repulsive hydration. J. Chem. Phys. 126, 044509 (2007).

67. Jusufi, A., Hynninen, A.-P. & Panagiotopoulos, A. Z. Implicit solvent models for micellization of ionic surfactants. J. Phys. Chem. B 112, 13783–13792 (2008).

68. Plimpton, S., Thompson, A. & Wood, M. LAMMPS as a tool in materials modeling workflows. in Proposed for presentation at the TMS 2020 Virtual Annual Meeting & Exhibition held March 15 - February 18, 2021 (US DOE, 2021). doi:10.2172/1853877.

69. Ham, F. S. Theory of diffusion-limited precipitation. J. Phys. Chem. Solids 6, 335–351 (1958).

