## Supplemental simulation parameters for "Interplay of condensation and chromatin binding underlies BRD4 targeting"

### Supplementary Information: Coarse-grained simulation parameters

In this section, we describe the coarse-grained force field and the simulation environment. Parameter values are provided in the tables below.

Non-bonded interactions between particles/blobs of types 1 and 2 are modeled via a combination of a repulsive WCA potential[S1] and an attractive Gaussian potential. For the WCA interactions, the length scale is $\sigma_{12} = (\sigma_1 + \sigma_2) / 2$, where $\sigma_1$ and $\sigma_2$ are the diameters of the hard cores of the particles, and the energy scale is set to 1 $kT$. For the Gaussian interactions between polymer blobs, the potential takes the functional form $U_{gauss}(r) = -A \exp(-Br^2)$ for $r < r_{cut}$, where the length-scale parameter is $B = 0.8 / [(R_{g,1} + R_{g,2}) / 2]^2$ and $R_{g,1}$ and $R_{g,2}$ are the radii of gyration of the blobs in dilute solution[S2]. The cutoff distance is set to $r_{cut} = 1.5\ (R_{g,1} + R_{g,2})$. The interaction-strength parameter, $A$, is chosen to be a constant value of -8 $kT$ for all interactions among N-terminal and C-terminal BRD4 blobs; however, because of the differing $\sigma$ and $R_g$ values of these species, the resulting well depths for the summed WCA and Gaussian potentials differ as well ($U_{min,NN}/kT$ = -0.75, $U_{min,CN}/kT$ = -1.2, $U_{min,CC}/kT$ = -1.9). These choices lead to the reproduction of the experimentally determined phase diagram as demonstrated in the main text. The interaction-strength parameter, $A$, between acetylated histone tails and the N-terminal BRD4 blob is varied to represent either weakly attractive interactions with $A$ = -25 $kT$ (resulting in a second virial coefficient for the blob-blob interactions of $b_2 = B_2/V_{endog}$ ~ -0.5, where $V_{endog}$ = ) or strongly attractive interactions with $A$ = -50 $kT$ (resulting in $b_2$ ~ -25).

**Table S1. Force-field parameters.**

| Parameter | Symbol | Value | Description |
|---|---|---|---|
| Occupied-volume diameter of the C-terminal blob (C) | $\sigma_C$ | 1 $\sigma$ = 4.94 nm | 678 disordered amino acids (AAs) |
| Occupied-volume diameter of the N-terminal blob (N) | $\sigma_N$ | 1.32 $\sigma$ | 361 disordered AAs + folded domains |
| Occupied-volume diameter of the Ferritin core (F) | $\sigma_F$ | 2.43 $\sigma$ | From Bracha et al. 2018[S3] |
| Occupied-volume diameter of the nucleosome (H) | $\sigma_H$ | 1 $\sigma$ | From Cutter and Hayes 2015[S4] |
| Occupied-volume diameter of the histone-tail blob (T) | $\sigma_T$ | 0.354 $\sigma$ | 30 disordered AAs |
| Radius of gyration of the C-terminal blob | $R_{g,C}$ | 0.812 $\sigma$ | Ideal polymer model for 678 disordered AAs |
| Radius of gyration of the N-terminal blob | $R_{g,N}$ | 0.845 $\sigma$ | Ideal polymer model for 361 disordered AAs and folded domains |

| | | | |
|---|---|---|---|
| Radius of gyration of the histone-tail blob | $R_{g,T}$ | 0.171 $\sigma$ | Ideal polymer model for 30 disordered AAs |
| FENE bond spring constant for N-C bonds | $K_{FENE,NC}$ | 2.886 $kT/\sigma^2$ | Ideal polymer model |
| FENE bond spring constant for F-C bonds | $K_{FENE,FC}$ | 1.937 $kT/\sigma^2$ | Ideal polymer model |
| FENE bond spring constant for H-H bonds | $K_{FENE,HH}$ | 5.0 $kT/\sigma^2$ | Ideal polymer model |
| FENE bond spring constant for T-H bonds | $K_{FENE,TH}$ | 5.704 $kT/\sigma^2$ | Ideal polymer model |

**Table S2. Simulation parameters for nucleation simulations.**

| **Parameter** | **Symbol** | **Value** | **Description** |
|---|---|---|---|
| Simulation temperature | $kT$ | 1 ε | Energy scale |
| Timestep for the molecular dynamics integration | $dt$ | 0.005 $\tau$ | |
| Langevin damping parameter | $\tau_{langevin}$ | 10 $\tau$ | >> velocity autocorrelation decorrelation time |
| Number of Corelets in Corelet nucleation simulations | $N_{corelet}$ | 512 | > 10 times the typical critical nucleus, $N^*$ |
| Number of BRD4 molecules in endogenous nucleation simulations | $N_{endog}$ | 5832 | > 40 $N^*$ |
| Size of a stable cluster of endogenous BRD4 molecules | $N_{nucl,endog}$ | 600 | > 4 $N^*$ |
| Size of a stable cluster of corelet molecules | $N_{nucl,corelet}$ | 200 | > 4 $N^*$ |
| Size of a minimal registerable cluster of endogenous BRD4 molecules | $N_{reg,endog}$ | ~43400 | $D_{reg}$ ~ 360 nm<br>$\rho_{condense}$ ~ 1.776e-3 $nm^{-3}$ |
| Size of a minimal registerable cluster of Corelet molecules | $N_{reg,corelet}$ | ~3070 | $D_{reg}$ ~ 360 nm<br>$\rho_{condense}$ ~ 1.256e-4 $nm^{-3}$ |

### Supplementary Information References

S1. Weeks, J. D., Chandler, D. & Andersen, H. C. Role of repulsive forces in determining the equilibrium structure of simple liquids. *J. Chem. Phys.* **54**, 5237–5247 (1971).

S2. Louis, A. A., Bolhuis, P. G., Hansen, J. P. & Meijer, E. J. Can polymer coils Be modeled as 'Soft colloids'? *Phys. Rev. Lett.* **85**, 2522–2525 (2000).

S3. Bracha, D. *et al.* Mapping Local and Global Liquid Phase Behavior in Living Cells Using Photo-Oligomerizable Seeds. *Cell* **176**, 407 (2019).

S4. Cutter, A. R. & Hayes, J. J. A brief review of nucleosome structure. *FEBS Lett.* **589**, 2914–2922 (2015).
